# Secreted retropepsin-like enzymes are essential for stress tolerance and biofilm formation in *Pseudomonas aeruginosa*

**DOI:** 10.1101/2025.03.18.643919

**Authors:** Justin D. Lormand, Charles H. Savelle, Jennifer K. Teschler, Eva López, Richard H. Little, Jacob G. Malone, Fitnat H. Yildiz, María J. García-García, Holger Sondermann

## Abstract

Proteases regulate important biological functions. Here we present the structural and functional characterization of three previously uncharacterized aspartic proteases in *Pseudomonas aeruginosa*. We show that these proteases have structural hallmarks of retropepsin peptidases and play redundant roles for cell survival under hypoosmotic stress conditions. Consequently, we named them retropepsin-like osmotic stress tolerance peptidases (Rlo). Our research shows that while Rlo proteases are homologous to RimB, an aspartic peptidase involved in rhizosphere colonization and plant infection, they contain N-terminal signal peptides and perform distinct biological functions. Mutants lacking all three secreted Rlo peptidases show defects in antibiotic resistance, biofilm formation, and cell morphology. These defects are rescued by mutations in the inactive transglutaminase transmembrane protein RloB and the cytoplasmic ATP-grasp protein RloC, two previously uncharacterized genes in the same operon as one of the Rlo proteases. These studies identify Rlo proteases and *rlo* operon products as critical factors in clinically relevant processes, making them appealing targets for therapeutic strategies against *Pseudomonas* infections.

**IMPORTANCE:** Bacterial infections have become harder to treat due to the ability of pathogens to adapt to different environments and the rise of antimicrobial resistance. This has led to longer illnesses, increased medical costs, and higher mortality rates. The opportunistic pathogen *Pseudomonas aeruginosa* is particularly problematic because of its inherent resistance to many antibiotics and its capacity to form biofilms, structures that allow bacteria to withstand hostile conditions. Our study uncovers a new class of retropepsin-like proteases in *P. aeruginosa* that are required for biofilm formation and bacterial survival upon stress conditions, including antibiotic exposure. By identifying critical factors that determine bacterial fitness and adaptability, our research lays the foundation for developing new therapeutic strategies against bacterial infections.

## INTRODUCTION

*Pseudomonas aeruginosa* is a Gram-negative bacterium frequently found in soil, aquatic settings and plants. Unfortunately, it is also an opportunistic pathogen in humans, causing acute and chronic life-threatening infections in immunocompromised and cystic fibrosis patients (1). Treatment of *Pseudomonas* infections is complicated by its intrinsic and acquired resistance to many antibiotics (2, 3), as well as by *P. aeruginosa’s* ability to form biofilms, multicellular communities embedded in a self-produced extracellular matrix (4). This extracellular matrix facilitates bacterial attachment to a variety of surfaces and provides a protective environment that allows bacteria to tolerate harsh environmental conditions, including attacks from the immune system or the presence of antimicrobials (5). Biofilms have been associated with the establishment of chronic infections (6). Also, biofilms facilitate the attachment of *P. aeruginosa* to implants, catheters and other medical devices, complicating their eradication from health care settings (2). Due to its multi-drug resistance and its threat for human health, the World Health Organization has identified *P. aeruginosa* as one of the critical priority ESKAPE pathogens (7). The development of effective therapies to combat *Pseudomonas* infections requires the identification of novel drug targets that aim to limit biofilm formation and bypass multi-drug resistance mechanisms.

Proteases, enzymes that catalyze the hydrolysis of peptide bonds, have been established as potent drug targets (8). Proteases can promote protein degradation into small peptides and amino acids, which can later be used for recycling or nutritional purposes, but they also constitute a major mechanism for the post-translational modification of proteins, providing venues for the regulation of their biological activities (9). Proteases have been classified on the basis of several criteria, including the location within the substrate where they hydrolyze peptide bonds (endopeptidases, exopeptidases, and omega-peptidases), their catalytic mechanism (serine, threonine, cysteine, aspartic and metalloproteases) and their evolutionary relationships (10, 11). To date, 2.8% of the proteins encoded in the *P. aeruginosa* genome have been annotated as enzymes with peptidase activity (12). Some of these peptidases have been characterized extensively for their roles in peptidoglycan homeostasis (13, 14). However, little is known about the biological roles of many others, particularly those belonging to the family of aspartic peptidases.

All aspartic peptidases are endopeptidases that use an aspartate dyad and an activated water molecule to hydrolyze the peptide bond (15). Some of the best-known aspartic peptidases belong to the AA clan, a superfamily with distinctive structural features, which includes pepsins and retropepsins (10). Pepsins (MEROPS family A1) are broadly distributed in eukaryotes and include enzymes with important roles in gastric digestion in chordates, while retropepsins (MEROPS family A2) were originally found in retroviral genomes, where they function to process polypeptide precursors into mature viral proteins (16). Although pepsins and retropepsins share little sequence identity, they have been proposed to be evolutionarily related as based on common structural features (17). Nonetheless, pepsins function as bilobed monomers, where each lobe contributes one of the aspartate residues of the catalytic dyad, while retropepsins are homodimers, with the aspartate dyad being formed at the dimer interface (18). For a long time, monomeric A1 pepsins were assumed to be restricted to eukaryotes, while dimeric A2 retropepsins were thought to be linked to retroviral activity. However, the massive sequencing of bacterial genomes has revealed that both families of aspartic peptidases are represented in prokaryotes as well (9, 19). Despite the widespread presence of AA clan peptidases in different bacterial species, only a few of them have been structurally or functionally characterized (9, 20–27).

In *Pseudomonas*, RimB is the only aspartic peptidase characterized to date. *P. fluorescens* RimB is co-transcribed as part of the *rim* operon, a locus involved in rhizosphere colonization and plant infection (26). RimB binds to and regulates the activity of RimK, an ATP-grasp fold α-L-glutamate ligase that catalyzes both the ATP-dependent synthesis of poly-α-L-glutamate peptides and the sequential addition of glutamate residues to the C-terminus of ribosomal protein RpsF (26, 28–30). RpsF poly-glutamylation by RimK has been shown to regulate ribosomal function and lead to proteome changes in response to environmental conditions (26). *In vitro* assays have shown that RimB can catalyze the proteolytic cleavage of glutamate residues on the C-terminus of RpsF, thereby counteracting RimK activity and dynamically regulating the levels of RpsF glutamylation (30). Additionally, RimB has been proposed to modulate cellular levels of glutamate/poly-α-L-glutamate (30).

Here we present the structural and functional characterization of three related, previously uncharacterized aspartic peptidases in *P. aeruginosa*. We show that these proteases have structural features characteristic of retropepsin-like peptidases, have sequence similarity to RimB and can cleave RimB substrates. However, they all contain N-terminal signal peptides suggesting that they localize in the periplasmic or extracellular space and have distinct roles compared to RimB. We show that mutants lacking all three secreted peptidases fail to thrive under hypoosmotic stress conditions. Based on these findings, we have named these proteases retropepsins linked to osmotic stress tolerance (Rlo). Our results provide evidence that Rlo proteases are structurally similar to retropepsins and have functions distinct to those of RimB, with requirements in osmotic stress tolerance, as well as in antibiotic resistance and biofilm formation.

## RESULTS

### Discovery of secreted aspartic proteases

A BLASTp search of the *P. aeruginosa* genome (strain UCBPP-PA14) identified three hypothetical proteins with sequence similarity to RimB. Based on our structural and functional characterization of these enzymes, we opted for calling them retropepsin-like osmotic stress tolerance peptidases (Rlo): RloA (PA14_41690), RloA2 (PA14_06040) and RloA3 (PA14_05970). A Clustal Omega alignment revealed that all three RloA proteins and RimB have a high degree of sequence identity around the N-terminally located DTG motif (Figure S1A), the characteristic active site consensus for clan AA aspartic proteases (31). Consistent with the presence of an aspartic protease active site, all RloA proteins are classified as aspartic peptidases in InterPro (IPR021109) (32). Overall, RloA proteins have about 31% sequence identity with RimB (Figure 1A-B, Figure S1B). However, a characteristic feature that sets RloA proteins apart from RimB is the presence of N-terminal signal peptide sequences (Figure S1A), suggesting that they are secreted and function in the periplasmic or extracellular space (33).

**Figure 1.**
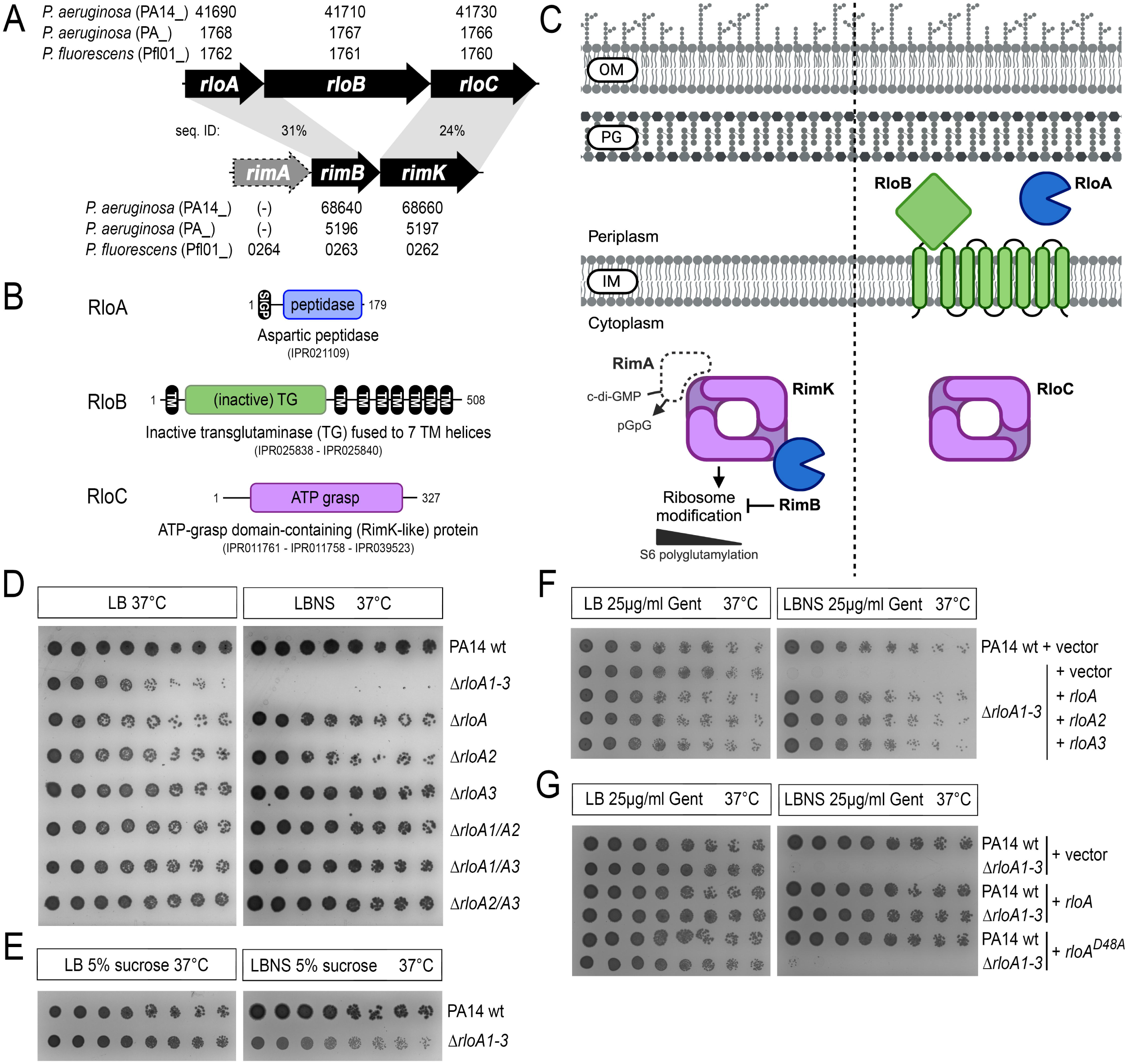
– RloA proteases and the *rlo* operon. **A.** Comparison between the *rlo* and *rim* operons. Genes are represented by arrows and their respective accession numbers are provided for each of the *Pseudomonas* species and strains indicated (PA14, PA01 and Pfl01). *rimA*, an EAL domain phosphodiesterase proposed to regulate RimK activity (26, 27) is shown with a grey discontinued line to highlight that this gene is only part of the *rim* locus in some *Pseudomonas* species. The percentages of sequence identity between RloA-RimB and between RloC-RimK are shown. In *P. aeruginosa*, there are two additional RloA homologs, RloA2 and RloA3 (not shown). Of these, RloA2 is predicted to be part of an operon together with genes encoding the CreBC two-component system. **B.** Domain organization of proteins encoded by the *rlo* operon. InterPro entries including each protein are indicated. **C.** Diagram comparing the subcellular localization of components encoded by the *rlo* and *rim* loci. The same shape and color were used to highlight similar proteins. OM, outer membrane. PG, peptidoglycan. IM, inner membrane. **D-G.** Titration assays showing growth of serial dilutions for the indicated bacterial strains in the media conditions shown at the top. LB, 1% agar Lysogeny Broth. LBNS, 1% agar LB without salts. In panels F and G, PA14 wild type (PA14wt) and Δ*rloA1-3* strains were transformed with either empty plasmids (vector) or plasmids encoding wild type or mutated versions of *rloA* proteases as indicated. Panel C was created with BioRender.com.

*rloA* is predicted to be part of an operon with two other uncharacterized genes that we have named *rloB* (PA14_41710) and *rloC* (PA14_41730) (Figure 1A). *rloB* encodes a hypothetical protein with an inactive transglutaminase domain fused to 7 transmembrane helices, while *rloC* encodes a hypothetical protein with an ATP-grasp domain (Figure 1B). The fact that both RloC and RimK contain an ATP-grasp domain, and that these proteins share 24% sequence identity, suggests some analogies between the *rlo* and *rim* loci (Figure 1A, C). Both genomic loci encode aspartic proteases (RimB and RloA) and the ATP-grasp proteins RloC and RimK are both predicted to localize to the cytosol (Figure 1C). However, a unique feature of the *rlo* operon is the presence of *rloB*, which encodes the only inactive transglutaminase of its kind in *Pseudomonas* genomes, a protein predicted to localize at the inner membrane, with the inactive transglutaminase domain exposed to the periplasmic space (Figure 1C). In summary, while two of the three proteins encoded in the *rlo* operon resemble the ribosome-modification proteins RimB and RimK, predictions of the sub-cellular localization and unique sequence features of the *rlo* gene products suggest a different biological function as those described for the RimK/RimB system.

Analysis of the phylogenetic distribution of the Rlo proteins across the tree of life revealed that RloA aspartic peptidases, classified as zinc proteases in Pfam (34), can be found mainly in proteobacteria (58% of the hits), as well as in members of Bacteroidota (20%) and Actinomycetota (7%) (Figure S2A). The most phylogenetically restricted protein is RloB, with 96% of the sequence hits coming from the phylum Proteobacteria. Conversely, ATP-grasp domain-containing proteins are widespread in bacteria and comprise a variety of protein architectures with distinct catalytic functions (35–37).

To investigate the evolutionary relationship between RimB and the RloA aspartic peptidases, we analyzed the taxonomic distribution of the RimB-RimK and RloA-RloB-RloC systems. This analysis revealed that the RimB-RimK system (Figure S2B, outer squares and circles) is more prevalent across bacteria than the *rlo* operon (Figure S2B, inner dark circles, triangles and squares) and that most organisms encoding an RloABC system also harbor RimB-RimK. Together, these findings suggest that the *rlo* operon might have originated from the *rim* locus and that, with the addition of the inactive transglutaminase encoded by *rloB* and the acquisition of signal peptide sequences by the *rloA* aspartic protease, the *rlo* locus acquired distinct functions evolutionarily advantageous for Gammaproteobacteria.

### RloA proteins are required for osmotic stress tolerance

In order to functionally characterize the biological roles of RloA proteases, we used allelic exchange in *P. aeruginosa* PA14 to generate deletion mutants in each *rloA*, *rloA2* and *rloA3*. None of these mutant strains showed any apparent defect in growth or behavior when compared to wild type PA14 (Figure 1D, rows 3-5). To explore whether the lack of defects in single *rloA* mutants could be due to functional redundancy amongst these proteases, we generated all possible combinations of double *rloA* mutants (Δ*rloA1/A2*, Δ*rloA1/A3* and Δ*rloA2/A3*) and a triple deletion of all RloA proteases (Δ*rloA1-3*). Mutant strains for double *rloA* mutants could be easily obtained and did not show any apparent defects (Figure 1D, rows 6-8). However, as we tried to obtain triple protease Δ*rloA1-3* mutants, we noticed that colonies surviving the second crossover counter-selection of the allelic exchange protocol (38) took a longer time to appear and were small-sized. Subsequent plating of these colonies on LB agar media failed to reproduce this phenotype, but PCR amplification and Sanger sequencing confirmed that these colonies were Δ*rloA1-3* mutants. Further analysis showed that Δ*rloA1-3* mutants had a normal growth pattern in liquid LB when compared with the parental wild type PA14 strain (Figure S3). Therefore, we suspected that the small colony size of Δ*rloA1-3* mutants during the allelic exchange selection process could have been due to effects of the hypoosmotic conditions of the second crossover counter-selection media (LB media without salts, supplemented with 15% sucrose). To test this hypothesis, Δ*rloA1-3* mutants were plated in LB media without salts (LBNS). We found that mutants lacking all three proteases failed to grow in LBNS media (Figure 1D, row 2). Moreover, this phenotype could be partially alleviated when sucrose, a known bacterial osmoprotectant (39), was added to the culture (Figure 1E), a result that is consistent with the growth of Δ*rloA1-3* mutants during the counter-selection step. Single and double mutants for *rloA* genes (Δ*rloA1*, Δ*rloA2*, Δ*rloA1/A2*, Δ*rloA1/A3* and Δ*rloA2/A3*) grew similar to the wild type strain on LBNS media (Figure 1D). Together, these results support that RloA, RloA2 and RloA3 are functionally redundant and uncover an important role for these novel secreted aspartic proteases in the mechanisms that allow *P. aeruginosa* to cope with osmotic stress.

We next aimed to rescue the lack of growth of Δ*rloA1-3* mutants in hypoosmotic media with plasmid-encoded RloA proteases. Ectopic expression of each individual RloA protease from these plasmids did not seem to affect growth in LB media. However, plasmid-encoded RloA proteases were able to rescue the growth of Δ*rloA1-3* mutants in LBNS (Figure 1F, rows 3-5). Plasmid-borne expression of *rloA*, *rloA2*, or *rloA3* resulted in similar levels of rescue, providing additional evidence that RloA proteases are functionally redundant.

To test whether the protease activity of RloA proteins is required for their roles in osmotic stress tolerance, we introduced an alanine mutation in the critical aspartic acid residue in the catalytic site of plasmid-encoded *rloA* (*rloA^D48A^*) and tested its ability to rescue the phenotype of triple protease Δ*rloA1-3* mutants in LBNS. We found that plasmid-encoded *rloA^D48A^* mutants were unable to rescue Δ*rloA1-3* mutant growth in hypoosmotic media (Figure 1G, rows 5-6). Therefore, RloA catalytic activity is required for its biological function regulating bacterial tolerance to osmotic stress.

### RloA proteases have retropepsin-like structural features

To better characterize the molecular features of RloA proteins, we determined the crystal structures of two representative orthologs, *P. fluorescens* RloA (RloA*_Pfl_*) and *P. aeruginosa* RloA2 (RloA2*_Pa_*), at a maximum resolution of 2.4 Å (Figure 2A; Table S1). RloA proteases fold into a β-sheet-rich structure, with a disordered internal segment (Figure 2A). Both proteins form C2 pseudosymmetric dimers in the crystal lattice. This dimeric state is consistent with their quaternary structure in solution as based on size-exclusion-coupled multi-angle light scattering (SEC-MALS) (Figure S4A). The conserved active site DTG sequence motif is located at the dimer interface of the proteases with the disordered flexible flap regions hovering above it (Figure 2A). Both RloA*_Pfl_* and RloA2*_Pa_* contain two cysteine residues that form a disulfide bond, stapling the distal tip of the β-strand succeeding the flexible region and the C-terminal tail together. These cysteine residues are conserved in periplasmic orthologs, but not in cytosolic orthologs such as RimB (Figure S1A, arrowheads).

**Figure 2.**
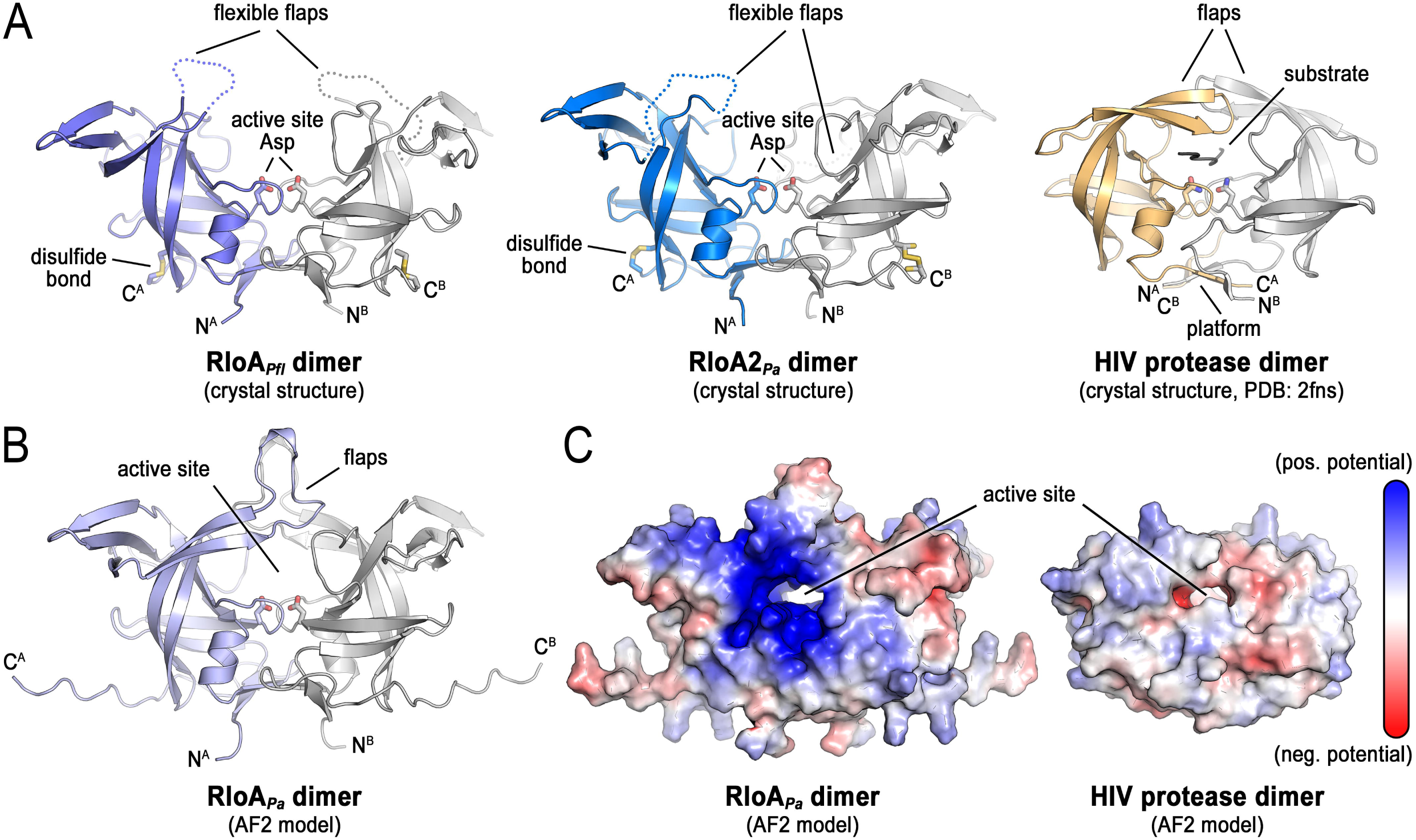
– Structural analysis of RloA proteases. **A.** Crystal structures of *P. fluorescens* RloA (left, violet-grey) and *P. aeruginosa* RloA2 (center, blue-grey) compared with the published HIV protease dimer structure (right, yellow-grey). The flexible flaps represent disordered regions corresponding to RloA*_Pfl_* residues 94-112 in chain A and 90-111 in chain B, and to RloA2*_Pa_* residues 137-159 in chain A and 141-152 in chain B. **B.** AlphaFold2 model of the *P. aeruginosa* RloA dimer. **C.** Surface views of the *P. aeruginosa* RloA and HIV protease AlphaFold2 dimer models illustrating residues with positive (blue) and negative (red) surface electrostatic potential.

A search against experimentally determined structures using the Foldseek server (40), with the crystallographic model of RloA*_Pfl_* as input, identified the following structurally similar proteins: an RloA-like protein from *Legionella pneumophila* (PDB: 2pma, sequence identity: 39.6%; Foldseek score: 515; Figure S4B); a yeast protein with a retroviral protease-like domain (sequence identity: 21.1%; Foldseek score: 117) (41); HIV protease (sequence identity: 14%; Foldseek score: 116; Figure 2A) (42); and retroviral-protease-like enzymes from *Leishmania major* and *Toxoplasma gondii* (sequence identity: 17 and 12%; Foldseek score: 108 and 107) (43, 44). The structural similarities between these proteins support the classification of RloA proteases as bacterial retropepsin-like enzymes. This classification is also supported by the fact that all three RloA proteases contain an alanine residue following the DTG motif (Figure S1A), a feature characteristic of retroviral aspartic proteases that differs from the serine or threonine residue common in pepsins (45).

A prominent feature from the structural comparison of RloA proteases with viral retropepsins is that the disordered regions in the RloA*_Pfl_* and RloA2*_Pa_* crystal structures correspond to the flaps that in HIV protease are described to close down on the substrate-bound active site in the holo conformation (46). Hence, the observed state in the RloA crystals likely depicts the apo conformation of the proteases. A model for the RloA*_Pa_*dimer obtained using AlphaFold2 (AF2) and the ColabFold server (47, 48) shows the flaps covering the active site through interactions of their distal tips (Figure 2B), forming a central channel for a substrate, analogous to the state described for substrate-bound HIV protease (49). However, the flaps of RloA*_Pa_* and its orthologs are longer than those of the HIV protease (Figures 2A and 2B), a feature that likely influences the accessibility and specificity of the active site to substrates.

RloA orthologs also differ from retroviral pepsins in the topology of their ‘β-sheet platform’ dimer interfaces. In HIV protease, the β-sheet platform comprises four interdigitated strands from the amino and carboxyl termini of the two protein protomers in the dimer (45). In contrast, the corresponding platform in the RloA*_Pfl_* and RloA2*_Pa_* dimers is formed without interdigitation of the strands, keeping the two protomers separate (Figure S4C, platform views). The respective platforms also differ in rotation relative to the dimer axis. Together, this organization is reminiscent of the topology observed in pepsins and in the eukaryotic retroviral-like protease Ddi1 (41). The β-sheet platform is presumed to stabilize the conformation of the enzyme active site (41). Therefore, variations in β-sheet platform organization may contribute to substrate specificity.

The active sites of viral and bacterial retropepsins have a similar topology, particularly regarding the placement of the conserved DTG motif that harbors the catalytic aspartate dyad. This observation suggests that a conserved reaction chemistry unifies these proteases (17). Additionally, both viral and bacterial retropepsins share a common two-fold rotational pseudosymmetry of the protease dimers, which likely contributes to substrate specificity, either by imposing symmetry constraints on the substrate or by perturbing the protease dimer upon substrate binding, as has been described for HIV protease (50). To gain initial insight into the potential nature of RloA substrates, we used a multiple-sequence alignment constructed with a representative set of RloA ortholog sequences and mapped the conservation onto the surface of the AF2 model of RloA*_Pa_* (Figure S4D). This analysis revealed considerable variability at the tips of the flaps and at the surfaces distal to the active site. In contrast, conservation was apparent at the active-site channel, indicating that RloA orthologs across different species have similar substrate preferences. We also found that, unlike the more hydrophobic environment observed for the HIV protease, the RloA protease creates a more polar, positively charged substrate environment (Figure 2C), suggesting that RloA substrates are negatively charged molecules.

In conclusion, the structural analysis of RloA proteases shows many common features with viral retropepsins, but also differences that might be relevant to define the accessibility of the active site and the substrate specificity of RloA proteases, such as variations in the length of the flexible flaps, the structure of the β-sheet platform, and the charge surrounding the active site channel.

### RloA cleaves glutamate-rich sequences

RimB, the cytoplasmic RloA homolog, has been shown to process polyglutamate residues (27, 30). Given that the structures of RloA orthologs indicate a positively charged substrate-binding interface (Figure 2C), we explored whether negatively charged polyglutamate peptides could be recognized and proteolytically cleaved by RloA proteases. For this, we first generated an AF2 model of RloA*_Pa_* dimers bound to a peptide with eight glutamate residues (Figure 3A). In this model, the peptide threads through a confined active site covered by the flaps. Each of the eight glutamate residues of the peptide engages in polar interactions with either lysine or arginine residues lining the active site of RloA*_Pa_* on either side. Pseudosymmetrical contacts involve residues K^46^, R^92^, R^114^, and R^137^ of RloA*_Pa_*. Except for R^92^, which is an alanine residue in RimB*_Pa_*, these positions are conserved in RloA paralogs and RimB (Figure S5A). Additional contacts with the substrate are asymmetric and involve K^95^, K^97^, and K^109^ from a single protomer of the RloA*_Pa_*dimer, residues that are not always conserved across bacterial retropepsin-like enzymes (Figure S5A). Also, the non-conserved residue R^99^, located in the flexible flaps, interacts with the backbone of the polyglutamate peptide (Figure 3A). The full coordination of the octaglutamate peptide in these models, along with interactions mediated by conserved residues in all RloA peptidases, indicates that RloA retropepsins could recognize and process polyglutamate (polyE) peptides.

**Figure 3.**
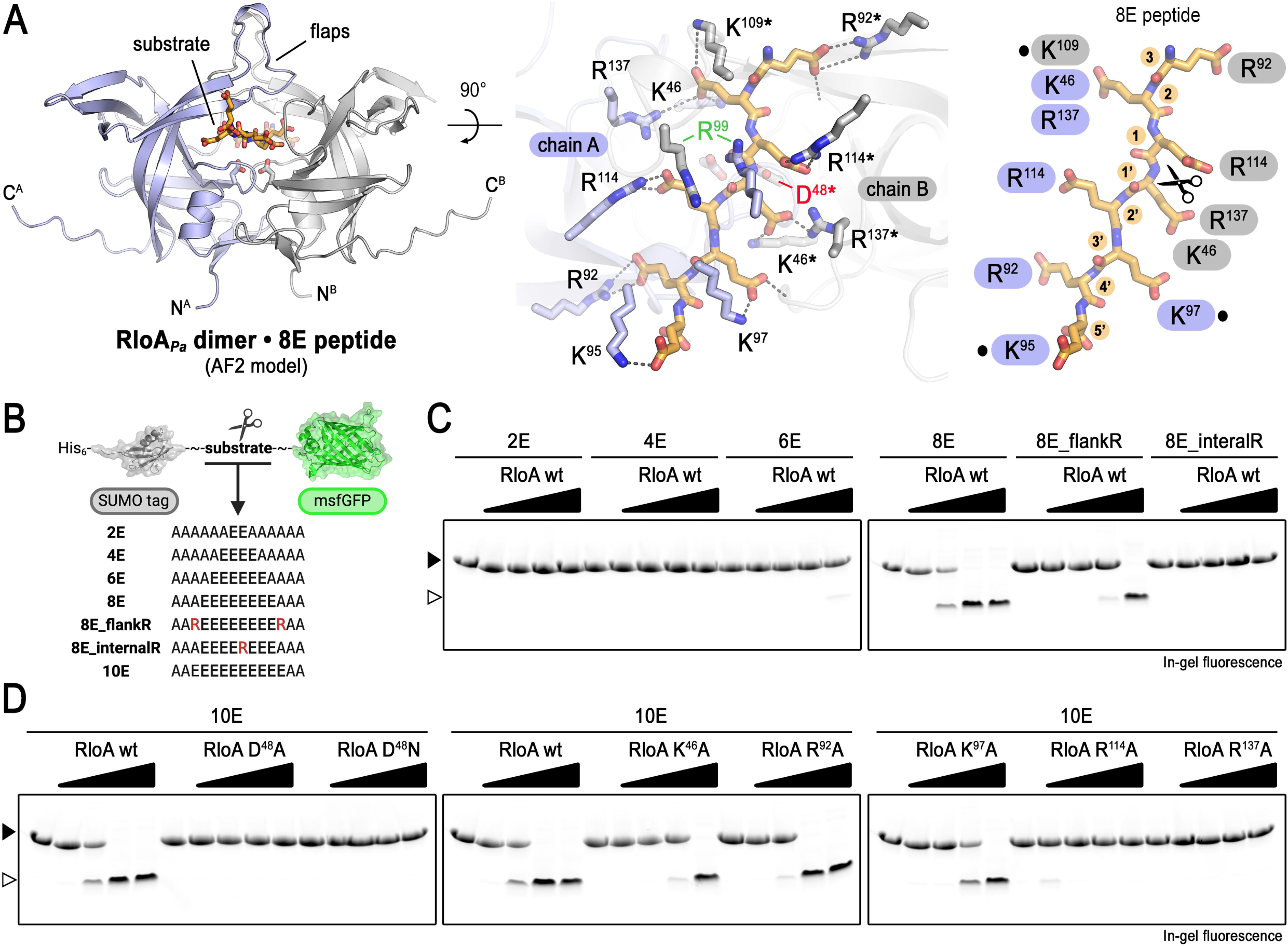
– Poly-glutamate peptides as RloA substrates. **A.** AlphaFold2 model of *P. aeruginosa* RloA dimer in complex with an 8 residue poly-glutamate peptide (orange). RloA protomers are highlighted in different colors (blue and grey). A close-up top view of the active site with the bound substrate is shown in the central panel. Residues involved in substrate coordination from each of the RloA protomers are highlighted in blue (chain A) and grey/asterisk (chain B). R^99^ residues involved in interactions with the backbone of the polyglutamate peptide are highlighted in green. The critical D^48^ residue at the conserved aspartic peptidase catalytic triad is highlighted in red. Right panel shows a schematic view of the substrate coordination interactions. Dots mark residues involved in asymmetric contacts with the substrate. Scissors highlight the scissile bond, which was predicted between the fourth and fifth residues of the substrate in the top-ranked model (additional registries, shifted by –1, were also observed in alternative models). **B.** Scheme illustrating pCleevR polyglutamate substrates. **C-D.** In-gel fluorescence after SDS-PAGE of the products from the enzymatic reactions between wild type or mutant RloA protease with different pCleevR constructs as indicated. Empty and filled arrowheads mark the positions of the cleaved and uncleaved substrates, respectively.

To test the structural prediction that polyglutamate could serve as a substrate for RloA orthologs, we used a reconstituted *in vitro* system to assess whether polyglutamated RpsF, a known substrate for RimB (27), could be processed by purified RloA*_Pa_*. As controls, we used RimB*_Pa_* and protease mutant versions in the critical active site aspartic acid residue previously shown to be required for catalytic activity in retropepsins (51, 52). We found that RloA*_Pa_* was able to cleave a genetically-modified RpsF containing eight consecutive glutamate residues at the C terminus (RpsF8E) as efficiently as RimB*_Pa_* (Figure S5B). Therefore, even though the cytoplasmic localization of RpsF makes it an unlikely substrate of RloA proteases, these experiments demonstrate that RloA is able to cleave polyglutamated proteins *in vitro*.

To further define the specificity of RloA proteases for polyglutamate substrates, we designed a fluorescent reporter that contained sequences with increasing number of glutamate residues (2E to 10E), flanked by a N-terminal SUMO and a C-terminal monomeric superfolder GFP (msfGFP) domain (Figure 3B, Figure S5C). We found that RloA was able to cleave reporters containing 8 or 10 consecutive glutamate residues, but failed to process substrates containing 6 or less glutamate residues (Figure 3C-D). Modifications of the 8E substrate introducing bulky arginine residues, either flanking the glutamates or in the middle of the 8E chain, respectively reduced or abrogated the catalytic efficiency of RloA (Figure 3C), indicating a strict sequence requirement for substrates at the catalytic site. Because the structural model predicts that conserved, positively charged arginine and lysine residues in RloA are responsible for polyglutamate coordination (Figure 3A), we tested whether amino acid substitutions in these residues affected the catalytic activity of RloA *in vitro* and/or the ability of RloA to rescue osmotic stress defects *in vivo*. As control, we used the catalytically inactive RloA^D48A^ and RloA^D48N^ mutants. We found that alanine substitutions at residues involved in polyE coordination decreased the catalytic activity of RloA, with residues closer to the catalytic dyad (R^114^A and R^137^A) having the strongest effects (Figure 3D). Additionally, mutations *rloA^R114A^* and *rloA^R137A^* failed to rescue the growth of Δ*rloA1-3* mutants in LBNS (Figure 4A). Together, our results validate the structural modelling of RloA bound to polyE peptides and demonstrate that RloA is capable of cleaving stretches of polyglutamate residues positioned both within proteins or at their C-termini.

**Figure 4.**
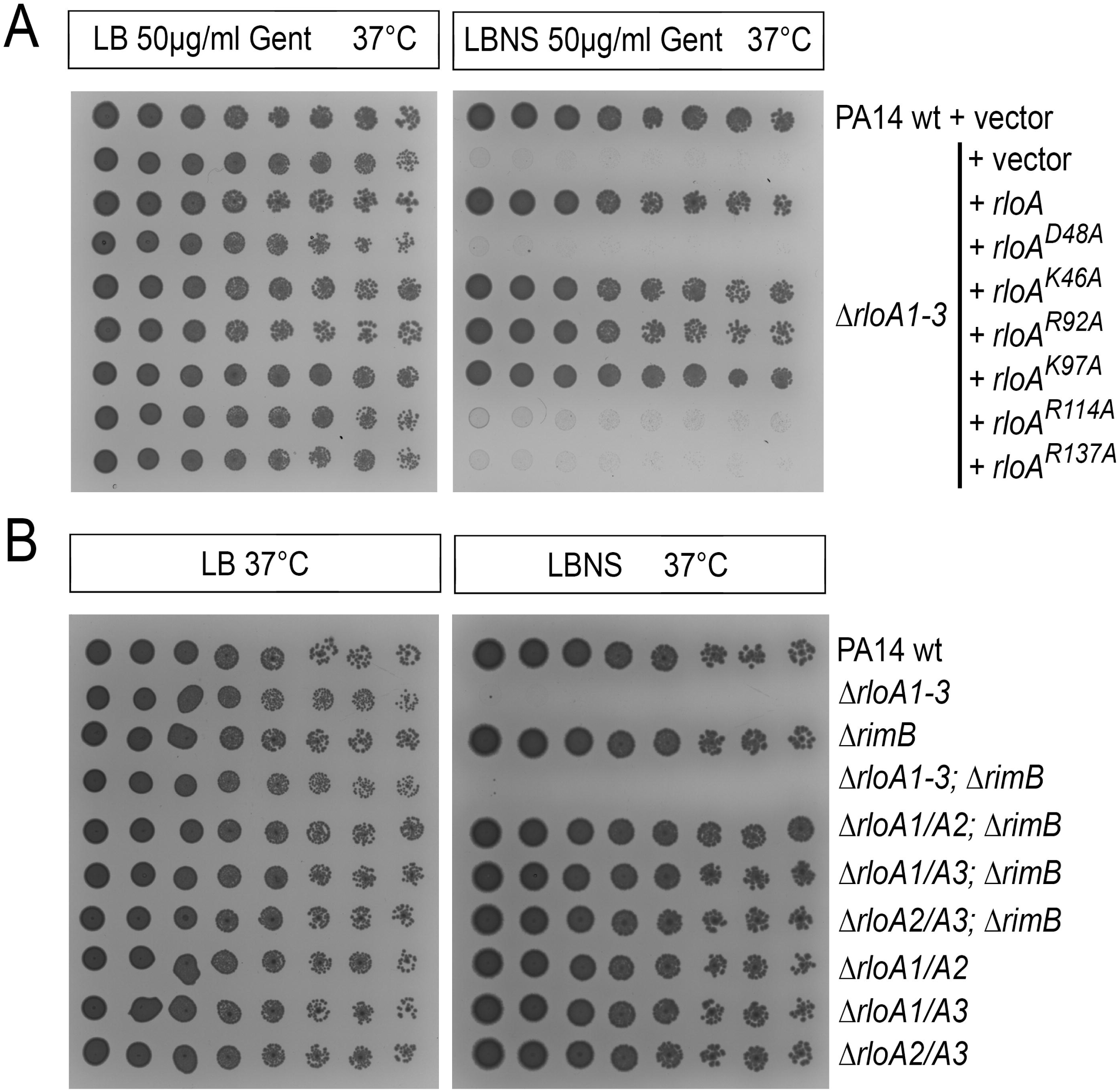
– Hypoosmotic stress tolerance experiments. Titration assays showing growth for serial dilutions of the indicated bacterial strains in the media conditions shown at the top. **A.** Resistance to osmotic stress was tested in PA14 wild type (PA14wt) and Δ*rloA1-3* strains transformed with either empty plasmids (vector) or plasmids encoding wild type or mutated versions of *rloA* proteases as indicated. **B.** Osmotic stress tolerance was tested for *rloA* and *rimB* mutant combinations. LB, 1% agar Lysogeny Broth. LBNS, 1% agar LB without salts.

In order to identify putative RloA *in vivo* substrates, we next queried the proteome of *P. aeruginosa* for genome-encoded proteins that localize in the periplasmic or extracellular space and contain 8 or more glutamate residues. This search did not identify any candidate. However, relaxing the criterium of our search to allow for proteins containing stretches of negatively charged residues (glutamate or aspartate) identified a single candidate: CprS (PA14_24340), a transmembrane histidine kinase featuring a DDDDEEEDDD sequence predicted to localize on a loop facing the periplasmic space (53). To test whether CprS could be cleaved by RloA, we designed a msfGFP fluorescent reporter containing a stretch of 19 amino acids from CprS that included the DDDDEEEDDD motif. When incubated with RloA, this reporter remained uncleaved (Figure S5D). Therefore, our findings suggest that genome-encoded proteins with stretches of negatively charged amino acids are not likely to be the physiological substrates of RloA proteases, but that RloA proteases rather cleave polyE peptides or post-translationally polyglutamated proteins.

Given the sequence and structural similarities between RloA proteases and RimB, as well as the fact that RloA can proteolytically cleave RimB substrates *in vitro* (Figure 3, Figure S5B), we wondered whether RimB could be functionally related to the biological roles of RloA proteases. To test this genetically, we generated Δ*rimB* deletion mutants in wild type *P. aeruginosa* PA14, as well as in PA14 strains carrying double mutant combinations of the RloA proteases (Δ*rloA1/A2,* Δ*rloA1/A3 and* Δ*rloA2/A3)* and the three RloA protease deletion mutant (Δ*rloA1-3)*. Compared to the corresponding parental strains, deletion of *rimB* did not alter growth in either LB media or hypoosmotic conditions for any of the mutant backgrounds tested (Figure 4B), a result that supports separate biological functions for RimB and RloA proteases.

### Deletion of RloA proteases affects biofilm formation and antibiotic sensitivity

We next asked whether deletion of the RloA proteases has other consequences for bacterial physiology, including effects on clinically relevant *Pseudomonas* processes such as biofilm formation and antibiotic resistance. We found that, compared to the wild type PA14 strain, Δ*rloA1-3* mutants failed to grow robust biofilms, as judged by the reduced amount of biomass visualized through crystal violet staining (Figure 5A). Additionally, Δ*rloA1-3* mutant colonies grown on Congo Red plates had a pale and smooth appearance (Figure 5B), consistent with a low production of extracellular matrix components (54). Mutants for individual (Δ*rloA1,* Δ*rloA2 and* Δ*rloA3* mutants) or double mutant combinations (Δ*rloA1/A2,* Δ*rloA1/A3 and* Δ*rloA2/A3)* of the RloA proteases showed a similar colony morphology to wild type PA14 on Congo Red plates (Figure S6B). However, Δ*rloA1* single mutants formed slightly less biofilm than the wild type strain when grown in microtiter plates (Figure S6A), albeit biofilm formation was not as disrupted as in the Δ*rloA1-3* strain. Notably, the inability of Δ*rloA1-3* mutants to grow biofilms could be rescued by plasmid-encoded RloA, but not by the *rloA^D48A^* catalytically inactive mutant version (Figure S7). Therefore, these results show that, in addition to their role helping cells cope with hypoosmotic stress, catalytically active RloA proteases are required for biofilm formation.

**Figure 5.**
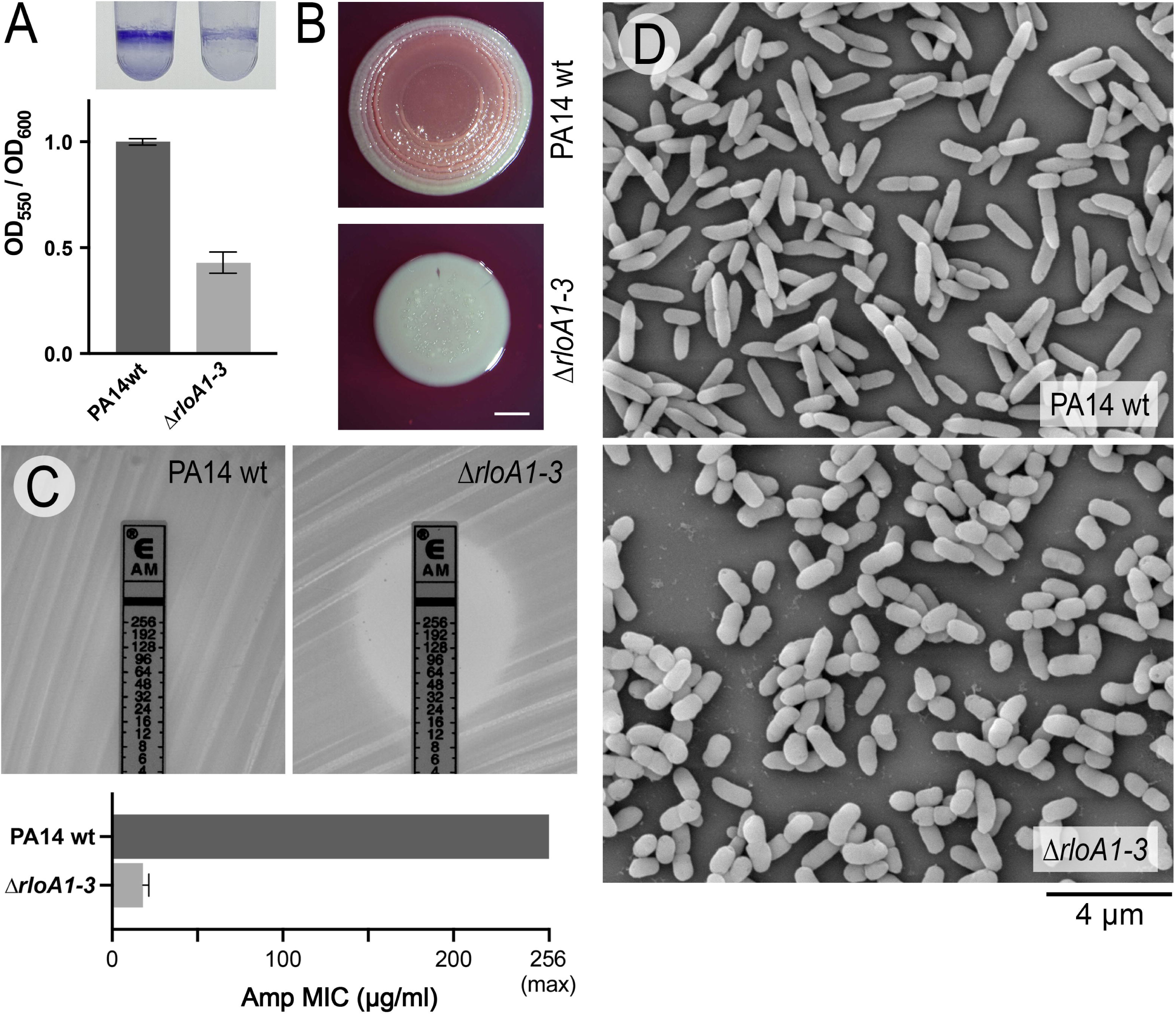
– Phenotypic characterization of RloA protease mutants in *Pseudomonas aeruginosa*. **A.** Biofilm formation in PA14 wild type and Δ*rloA1-3* mutants was assessed through crystal violet staining of overnight cultures grown in 0.4% Arginine M63 media (top). Graph shows the normalized average intensity of crystal violet staining (OD_550_) relative to the growth of the corresponding bacterial culture (OD_600_), n=4. Error bars show standard deviation. **B.** PA14 wild type and Δ*rloA1-3* mutant colonies grown in Congo Red 0.2% Glucose M63 agar. Scale bar represents 2 mm. **C.** Ampicillin ETEST strips tested on lawns of PA14 wild type and Δ*rloA1-3* mutant strains grown on Mueller-Hinton agar. Graph shows average Minimal Inhibitory Concentration (MIC) from triplicate experiments. Error bars show standard deviation. **D.** Scanning electron micrographs of PA14 wild type (top) and Δ*rloA1-3* mutant cells (bottom). Scale bar represents 4 μm.

To test whether RloA proteases influence antibiotic resistance, we first analyzed our PA14 wild type and Δ*rloA1-3* mutant strains through a comprehensive panel of clinically relevant antibiotic conditions. This analysis revealed major differences in susceptibility between wild type PA14 and Δ*rloA1-3* mutants for antibiotics that inhibit cell wall synthesis. Specifically, Δ*rloA1-3* mutants showed increased sensitivity to β-lactam antibiotics ampicillin, amoxicillin, and cephalexin (Figure S8). In contrast, Δ*rloA1-3* mutants showed either minor or no differences in sensitivity with respect to the wild type PA14 strain when exposed to antibiotics that inhibit ribosomal function or nucleic acid synthesis (Figure S8). The increased sensitivity of Δ*rloA1-3* mutants to β-lactam antibiotics was further confirmed using ampicillin ETEST strips (Figure 5C, Figure S6C). *P. aeruginosa* resistance to most β-lactam antibiotics is mediated by multidrug efflux systems, which pump antimicrobials out of the cell, and by the action of β-lactamases, enzymes that specifically degrade β-lactam antibiotics (3, 55). We found that Δ*rloA1-3* mutants showed increased sensitivity to augmentin, a drug that combines a β-lactam antibiotic with the β-lactamase inhibitor clavulanic acid (Figure S8). Therefore, it is unlikely that the differences in antibiotic sensitivity between PA14 wild type and Δ*rloA1-3* mutants are due to variations in the levels of β-lactamase activity. It is possible that RloA proteases influence either the permeability of the outer membrane to β-lactam antibiotics (through regulating in the activity of porin channels) or the function of efflux pumps. However, porins and efflux pumps frequently mediate resistance to multiple antimicrobials with diverse modes of action (3, 55). Given that major differences in sensitivity are only observed for β-lactam antibiotics, our results argue that the basis for these differences might relate to the common mechanism of action of these antimicrobials on cell wall integrity. β-lactam antibiotics inhibit the transpeptidase activity of penicillin-binding proteins (PBPs), generating an imbalance between peptidoglycan synthesis and remodeling processes that ultimately results in periplasmic stress and toxicity (56). Therefore, it is possible that the absence of RloA proteases causes periplasmic stress and/or defects in cell wall homeostasis that predispose Δ*rloA1-3* mutants for a higher sensitivity to β-lactam antibiotics.

### RloA proteases regulate cell morphology

The ability of bacteria to tolerate osmotic changes in the environment partially depends on the integrity of their cell wall, a peptidoglycan structure that encases the cytoplasmic membrane, providing strength to resist turgor pressure and regulating bacterial shape (57). To explore whether the inability of triple protease Δ*rloA1-3* mutants to grow under hypoosmotic conditions and their higher sensitivity to β-lactam antibiotics could be caused by defects in cell wall homeostasis, we asked whether cell morphology is affected by deletion of RloA proteases. Both phase contrast and scanning electron microscopy (SEM) revealed that Δ*rloA1-3* mutants exponentially growing in LB showed an abnormal cell shape compared to the wild type PA14 strain (Figure 5D, Figure S9). Wild type PA14 cells had the typical rod-shaped morphology characteristic of *Pseudomonas* bacilli, with an average length of 2 μm and an average width of 0.64 μm. In contrast, Δ*rloA1-3* mutant cells were significantly shorter (average 1.5 μm) and wider (average 0.8 μm), giving them a chubby appearance. Deletion of individual RloA proteases (Δ*rloA1*, Δ*rloA2* and Δ*rloA3* mutants) or double protease mutant combinations (Δ*rloA1/A2*, Δ*rloA1/A2* and Δ*rloA2/A3* mutants) did not cause major alterations in either the length or width of bacteria (Figure S9). Therefore, these results indicate that RloA proteases have redundant roles in the control of cell morphology. The severe morphological defects of Δ*rloA1-3* mutants suggest a possible imbalance between the mechanisms that coordinate cell elongation, by lateral expansion of the peptidoglycan layer, and those that promote the synthesis of peptidoglycan at the septation ring during cell division (58).

### The *rlo* operon controls osmotic stress tolerance, biofilm formation and antibiotic resistance

Because genes co-transcribed as part of an operon often have related functional roles (59), we wondered if RloB and RloC control the same cellular processes as RloA proteases. To test this, we used allelic exchange to generate mutations in *rloB*, *rloC* and the complete *rlo* operon (Δ*rloABC*), as well as mutant combinations involving the three *rloA* proteases, *rloB* and *rloC*. Interestingly, we found that deletion of *rloB*, *rloC*, or both in a triple protease Δ*rloA1-3* mutant background (Δ*rloA1-3;* Δ*rloB,* Δ*rloA1-3;* Δ*rloC* and Δ*rloA1-3;* Δ*rloBC* mutants) completely rescued the inability of Δ*rloA1-3* mutants to grow under hyposmotic stress conditions (Figure 6A) and partially rescued their antibiotic resistance, biofilm formation and cell morphology phenotypes (Figure 6B-J). Individual Δ*rloB* and Δ*rloC* mutants showed an increased sensitivity to ampicillin compared to wild type (Figure 6C-D), as well as cell morphology defects (Figure 6E-H). Also, Δ*rloC* mutants formed less biofilm compared to wild type cells (Figure 6B). However, Δ*rloB* and Δ*rloC* mutant defects in these processes were not as pronounced as those of triple protease Δ*rloA1-3* mutants. For instance, Δ*rloB* and Δ*rloC* cells were shorter, but their width was comparable to that of wild type cells (Figure 6E-H). Altogether, our genetic analysis is consistent with the lack of *rloB* and *rloC* function being epistatic to the defects of Δ*rloA1-3* mutants, providing genetic evidence that RloA proteases and the products of the *rlo* operon play roles in the same molecular processes that ultimately regulate hypoosmotic sensitivity, biofilm formation, antibiotic resistance and cell morphology.

**Figure 6.**
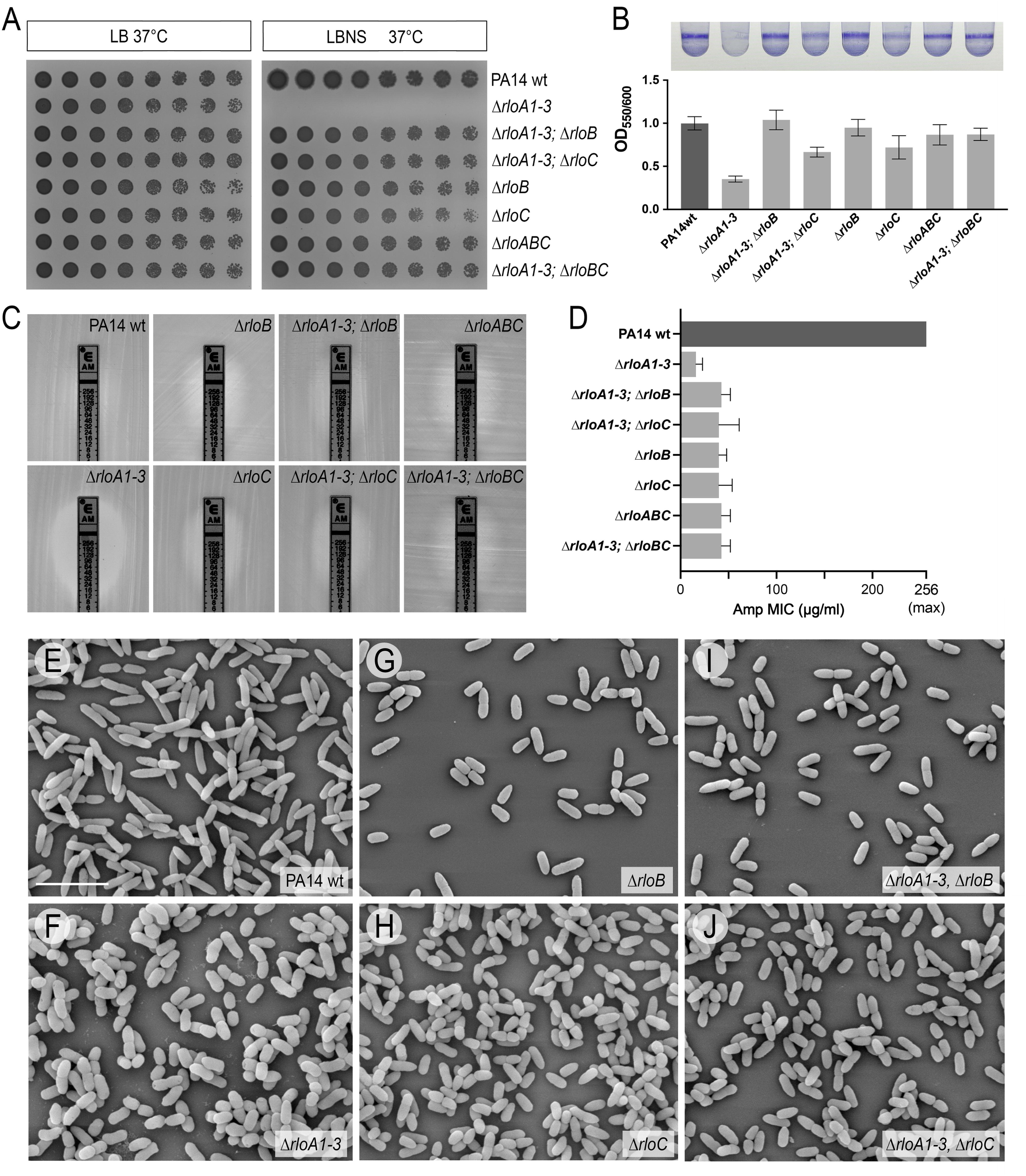
– Phenotypic characterization *rlo* operon mutants. **A.** Growth of serial dilutions for the bacterial strains indicated at each row in the media conditions shown at the top. LB, 1% agar Lysogeny Broth. LBNS, 1% agar LB without salts. **B.** Biofilm formation in PA14 wild type and indicated mutants was assessed through crystal violet staining of overnight cultures grown in 0.4% Arginine M63 media (top). Graph shows the normalized average intensity of crystal violet staining (OD_550_) relative to the growth of the corresponding bacterial culture (OD_600_), n=4. Error bars show standard deviation. **C.** Representative results from ampicillin ETEST strips as tested on lawns of PA14 wild type and indicated mutant strains grown on Mueller-Hinton agar. **D.** Graph shows average Minimal Inhibitory Concentration (MIC) from triplicate ampicillin ETEST strip experiments. Error bars show standard deviation. **E-J.** Scanning electron micrographs of PA14 wild type cells and mutants of the indicated genotypes. Scale bar in represents 4 μm.

## DISCUSSION

Our structural and phenotypic characterization of aspartic proteases in *Pseudomonas aeruginosa* uncovered three previously uncharacterized enzymes with important roles in hypoosmotic stress tolerance, antibiotic resistance, and biofilm formation.

We show that RloA proteins have structural features characteristic of retroviral pepsins, including a C2 pseudosymmetric dimeric arrangement, the location of the active site at the dimer interface, and the presence of flexible flaps covering the substrate channel. Studies on HIV protease have determined that substrate recognition by retroviral pepsins is dictated by the fitting of the substrate shape and volume at the substrate cavity, rather than by the recognition of a specific amino acid sequence in the substrate (60). In this sense, we identified crucial differences between HIV protease and RloA proteases at motifs previously established to play important roles in substrate accessibility. These include the presence of longer flexible flaps, which in HIV protease undergo major conformational changes between the apo and holo states (49) and a different organization of the β-sheet platform, which has been proposed to stabilize the conformation of the active site (41). Additionally, we found that, in contrast to the hydrophobic environment of the HIV protease active site, RloA proteases have a positively charged substrate channel, a feature that likely determines an affinity for negatively charged substrates.

The three RloA proteases described in our study are related to RimB, the only aspartic protease previously characterized in *Pseudomonas* (26, 27). In addition to a high degree of sequence conservation at the active site, RloA is part of an operon that bears some parallels with the RimB-RimK system (Figure 1A, C) and we show that RloA can cleave RimB substrates *in vitro* (Figure 3, Figure S5B). Despite these analogies, the presence of a signal peptide in RloA proteases predicts a different subcellular localization to RimB. Moreover, we found that deletion of *rimB* does not genetically interact with combinations of RloA protease mutants (Figure 4B). Therefore, while both RimB and RloA can recognize and cleave polyglutamate substrates *in vitro*, our data supports that these proteins evolved to perform different biological functions in distinct compartments.

Our structural models and *in vitro* experiments indicate that RloA proteases can cleave polyglutamate substrates. The fact that residues predicted to participate in glutamate coordination are conserved across RloA proteases (Figure S5A), coupled with the observation that mutations in these residues impair the ability of RloA to cleave polyglutamate *in vitro* (Figure 3D) and to complement the phenotype of Δ*rloA1-3* mutants (Figure 4A), collectively suggest that the cleavage of polyglutamate is important for the *in vivo* functions of RloA proteases. Bioinformatic searches failed to identify candidates containing polyglutamate stretches. Therefore, RloA physiological substrates are unlikely to be genome-encoded proteins. It is possible that, similar to RimB, RloA proteins cleave substrates that become polyglutamated post-translationally. Polyglutamylation is a reversible post-translational modification first discovered on tubulins and recently reported to regulate a few other eukaryotic proteins (61–63). In bacteria, the only known proteins to undergo polyglutamylation are RpsF and the *Legionella* ubiquitin ligase SdeA (26, 28, 64), none of which are plausible substrate candidates for RloA proteases. Another possibility is that RloA cleaves free polyglutamate peptides. Poly-γ-glutamate, a natural polymer produced by certain Gram-positive bacteria, is present as either a surface-bound or extracellular polypeptide and has been involved in protecting bacteria from environmental insults and/or as a source of glutamate during stationary phase starvation (65). Although *Pseudomonas* cannot synthesize poly-γ-glutamate, poly-α-glutamate polymers might have similar roles. In this respect, it is interesting to note that glutamate can function as an osmolyte, helping bacteria cope with osmotic stress (66). Additional experiments will be required to address whether RloA proteases regulate polyglutamylated proteins and/or glutamate-polyglutamate homeostasis.

Transposon insertions in *rloA* had been found in several suppressor screens related to extracellular matrix production and antibiotic resistance (67–69). However, the specific roles of RloA proteases in these processes had not been characterized. Here, we provide evidence that RloA peptidases have redundant roles to help cells cope with hypoosmotic stress conditions, as well as with the ability of *P. aeruginosa* to form biofilms and to survive in the presence of antibiotics that target the cell wall. At present, it is unclear whether all Δ*rloA1-3* phenotypes stem from requirements in a common molecular mechanism or if RloA proteases could influence several cellular processes. In this regard, it is worth noting that cell shape defects in Δ*rloA1-3* mutants are observed in the absence of any environmental stressors and therefore are likely a primary consequence of RloA loss of function. Since the peptidoglycan cell wall regulates cell shape and provides strength to resist turgor pressure (57) it is possible that the inability of Δ*rloA1-3* mutants to survive in hypoosmotic media is linked to the abnormal morphology we observed in Δ*rloA1-3* cells and due to defects in peptidoglycan homeostasis. Peptidoglycan defects could also explain the higher sensitivity of Δ*rloA1-3* mutants to antibiotics that specifically inhibit peptidoglycan synthesis. Also, established links between peptidoglycan homeostasis and biofilm formation (70–72) suggest that the failure of Δ*rloA1-3* mutants to grow biofilms could also originate from cell wall defects. Therefore, it is tempting to hypothesize that RloA peptidases could directly or indirectly regulate the activity of peptidoglycan synthases or hydrolases (58, 73). Regardless of whether the different phenotypes of Δ*rloA1-3* mutants stem from defects in one or multiple molecular processes, RloA proteases emerge as attractive molecular targets to fight *Pseudomonas* infections, given the clinical importance of addressing both antibiotic resistance and biofilm formation in these species.

Our research also uncovered important requirements for *rloB* and *rloC*, the other two genes encoded by the *rlo* operon. Our finding that mutations in *rloB* and *rloC* can rescue the defects of Δ*rloA1-3* mutants provides genetic evidence that all genes in the *rlo* operon control the same physiological processes. Mutations in *rloB* and *rloC* have been found in a number of genetic screens and experimental evolution experiments linked to antibiotic susceptibility and virulence (67–69, 74–80). Additionally, a number of Tn-seq studies have identified links between the *rlo* operon and *rloA* proteases in host colonization, virulence and/or switch to a chronic infection state (81–85). Interestingly, mutations in *rloB* and *rloC* are consistently found in these Tn-seq studies to have the most drastic effects on bacterial survival, while individual mutations in *rloA* proteases are only found to have moderate, context-specific or no effects on bacterial fitness. This lack of consistent fitness trade-offs in individual *rloA* mutants is consistent with our results showing the functional redundancy of *rloA* proteases and illustrates the limitations of genetic screens to identify genes with redundant functions.

Although the *rlo* operon is widely conserved in Gammaproteobacteria, little is known regarding the roles of this locus in other bacteria. In *Vibrio cholerae*, *rloABC* orthologous genes are amongst the most upregulated loci controlled by VxrB, the response regulator of the VxrAB two-component signaling system, which is linked to intestinal colonization, biofilm formation, type 6 secretion and cell wall integrity (86–88). However, the functional role of the *V. cholerae rlo* operon in these processes remains to be explored.

While additional experiments will be required to unravel how the protein activities of RloA, RloB and RloC ultimately influence the physiological processes controlled by these enzymes, our findings identify RloA proteins as a novel class of secreted bacterial retropepsin-like enzymes and uncover important roles for these peptidases, as well as for the other genes encoded in the *rlo* operon, in hypoosmotic stress tolerance, antibiotic resistance, biofilm formation, and cell morphology.

## MATERIALS AND METHODS

### Bacterial Strains, Plasmids, Growth Media and Culture Conditions

Cells were grown in Lysogeny Broth (LB) growth media, LB media without salts (LBNS), or Terrific Broth (TB) at 37°C or 18°C as indicated. Details about strains used and antibiotic selection can be found in Tables S2-S3.

*Pseudomonas aeruginosa* UCBPP-PA14 strain was genetically modified using allelic exchange (38). Targeting constructs were delivered by conjugation with S17.1 λ*pir E. coli* cells. Engineered strains and targeting vectors are listed in Supplementary Tables. Complementation and plasmid-encoded expression of genes in *P. aeruginosa* was performed using electroporation (89, 90).

For titration assays, overnight cultures grown at 37°C were diluted to OD_600_ 0.1, then 10-fold serial dilutions were prepared in LB and 3 μl of each dilution were spotted onto 1% agar LB or LBNS plates. Plates were grown overnight at 37°C and imaged using a BioRad ChemiDoc MP imager.

For assessment of extracellular matrix formation in colony biofilms, overnight bacterial cultures were diluted to OD_600_ 0.1, then 3 µl were spotted onto Congo Red plates (1.5% agar M63 minimal media supplemented with 1mM MgSO_4_, 40 µg/ml Congo Red, 20 µg/ml Coomassie Blue and either 0.5% casamino acids and 0.2% glucose, or 0.4% arginine). Plates were grown at 37°C overnight, then at room temperature (16-20°C) for 6 days inside a 99% humidity chamber.

For growth curves, overnight cultures were diluted to OD_600_ 0.1 in LB or LBNS media. Bacterial dilutions (100 µl) were aliquoted in 96-well polystyrene plates at least in triplicate. Plates were sealed with parafilm and grown at 37°C with shaking in a Tecan Infinite 200Pro plate reader. OD_600_ was measured every 20 minutes for 12-18 hours.

For biofilm assays, overnight cultures were diluted to OD_600_ 0.1 into M63 minimal media supplemented with 1 mM MgSO_4_ and 0.4% arginine or with 1 mM MgSO_4_, 0.5% casamino acids and 0.2% arabinose. Bacterial dilutions (100 μl) were distributed into 96-well PVC plates. Plates were sealed with parafilm and incubated at 37°C for 12-16 hours without shaking. Biofilms were stained with crystal violet (91). Biofilms were quantified as the relative OD_550_/OD_600_ normalized to the wild type average.

For ampicillin sensitivity assays, overnight cultures were diluted to OD_600_ 0.1 on LB, then swabbed 3 times onto Müller-Hinton Agar plates using a cotton applicator. Ampicillin ETEST strips (ETEST AM 0.016-256 μg/mL) were applied 5 minutes afterwards and inhibitory halos were quantified after incubation at 35°C for 16-18 h.

### Cloning and site-directed mutagenesis

PCR amplified or gene synthesized fragments were cloned using InFusion (Takara) or SLiCE (92) cloning. Overlap extension PCR was used to generate deletion alleles for allelic exchange. For site-directed mutagenesis, QuikChange II (Agilent) or SPRINP (93) were used. Primers are listed in Table S4. All constructs were confirmed by Sanger DNA sequencing.

### Bioinformatics and software

Conserved domains were identified using InterPro and Pfam databases (32, 34). Phylogenetic distribution of the relevant protein families was plotted using Annotree (94). EggNOG version 4.5.1 (95) was used to identify orthologous proteins to RimK, RimB, RloA1-3, RloB and RloC. The final list of representatives was curated manually and illustrated with iTol (96). Subcellular localization predictions were obtained from Pseudomonas.com (12) and PSORTb (33). SignalP was used to predict signal peptide sequences (97). ConSurf was used to map sequence conservation onto protein structures (98). Sequence logos were generated with WebLogo (99). Geneious Prime software was used for Sanger sequence analysis and Clustal Omega alignments. Primer3 was used for primer design (100). Prism software (GraphPad) was used for data plots and for statistical analysis using paired two-tailed parametric t-tests (p values higher than 0.05 were not considered statistically significant).

### Protein expression and purification

*E. coli* BL21 T7 Express cells (New England Biolabs) transformed with His_6_-SUMO-pET28a, pCleevR or pETM11-derived plasmids were grown in Terrific Broth (TB) with appropriate antibiotics at 37°C. Protein expression was induced with 0.5 mM IPTG when cultures reached OD_600_ ∼1, then cells were cultured for 16 hours at 18 °C. Cells were lysed through freeze-thawing and sonication. Cleared cell lysates were incubated with Ni-NTA Superflow resin (Qiagen) for 1 hour in Ni-NTA binding buffer (25 mM Tris-HCl pH 8.5, 500 mM NaCl, 20 mM imidazole), then washed with 30 column volumes of Ni-NTA binding buffer by gravity flow. Proteins were eluted in 6 column volumes of Ni-NTA elution buffer (25 mM Tris-HCl pH 8.5, 500 mM NaCl, 400 mM imidazole). Eluates were buffer exchanged into gel filtration buffer (25 mM Tris-HCl pH 7.5, and either 300 mM NaCl [or 150 mM NaCl for PA0462]) via a HiPrep 26/10 desalting column (GE Healthcare). RloA His_6_-SUMO-tagged proteins were incubated overnight with Ulp-1-His_6_ protease to cleave the tags and then separated from His_6_-SUMO and ULP1-His_6_ as the flow through from a HisTrap Ni-NTA column (GE Healthcare). Proteins were concentrated with an Amicon Ultra 10K concentrator to 3 ml prior to size-exclusion chromatography onto a HiLoad 16/60 Superdex 200 gel filtration column (GE Healthcare) equilibrated in gel filtration buffer. Fractions containing purified protein were identified by SDS-PAGE, concentrated, frozen in liquid nitrogen, and stored at –80°C.

### Crystallography

Dilutions of each protein in the appropriate gel filtration buffer were equilibrated to crystallization temperature. Crystals were grown at 20°C using hanging drop vapor diffusion with 1:1 mixture of protein to reservoir buffer (drop size 1.6 µl). Crystals formed with the following conditions: 10 mg/ml PA0462 (18–169) in 0.25 M sodium phosphate dibasic and 24% PEG 3350; or 5 mg/ml Pfl01_1762 (20–178) in 0.2M ammonium acetate, 20% PEG 3350 and 0.1 M HEPES pH 8.0. Crystals were equilibrated for at least 7 days before harvesting. Crystals were cryoprotected in reservoir buffer with 20% xylitol, then flash frozen and stored in liquid nitrogen. Data were collected by synchrotron radiation at Cornell High Energy Synchrotron Source (CHESS) and NE-CAT 24ID-C and 24ID-E beamlines at the Advanced Photon Source (APS) at Argonne National Laboratory. Diffraction data sets were processed using XDS, Pointless and Scala (101, 102). The initial structures of Pfl01_1762 and PA0462 were solved by molecular replacement using Phenix (103) and the unpublished coordinates of *Legionella pneumophila* protein Lpg0085 (PDB:2pma) as the search model. Building and refinement were carried out with Coot (104) and Phenix (103). Illustrations were prepared with the Pymol Molecular Graphics System, version 2.5.7 (Schrödinger, LLC). Electrostatic potential maps were calculated using the Pymol plugin for APBS (105). All software packages were accessed through SBGrid (106).

### Size-exclusion chromatography-coupled multi-angle light scattering (SEC-MALS)

5 mg/ml (289 µM) of purified PA0462 (18–169) or PA1768 (23–179) protein was injected onto a Superdex 200 Increase 10/300 gel filtration column (GE Healthcare) equilibrated with gel filtration buffer (25 mM Tris-HCl pH 7.5, 300 mM NaCl). An Agilent G1310A isocratic pump was used to run samples at a flow rate of 0.7 ml/min through the gel filtration column coupled to a static 18-angle light scattering detector (DAWN HELEOS-II, Wyatt Technology) and a refractive index detector (Optilab T-rEX, Wyatt Technology). Data were collected every second. Data analysis was performed with Astra 6.1 (Wyatt Technology) yielding the molar mass and mass distribution (polydispersity) of the sample. Monomeric BSA (Sigma; 5 mg/ml) was used to normalize the light scattering detectors.

### Enzymatic assays

Purified RpsF or RpsF8E (5 µM final) were incubated with RimB, RloA, or their mutant variants (0.1 µM final) in reaction buffer (25 mM HEPES-NaOH pH 7.5, 150 mM NaCl) for 2 hours at 25°C. Reactions were subjected to SDS-PAGE using 4-10% gradient gels (Bio-Rad), followed by Coomassie staining. pCleevR substrates (30 µM final) were incubated with RloA wild type or mutant variants (10x serial dilutions; range: 0.3 nM to 300 nM) for 30 min at 25°C. Reactions were stopped by addition of SDS-PAGE sample buffer and un-boiled samples were analyzed by SDS-PAGE.

### Imaging

Overnight cultures of the indicated strains were diluted 1:100 on LB media and grown at 37°C to OD_600_ 0.4-0.6. For SEM, Cells were washed in phosphate buffered saline (PBS) and fixed overnight at 4°C in a 2.5% glutaraldehyde solution placed on top of poly-D-lysine coated coverslips. Coverslips were washed in PBS, fixed in 1% OsO_4_ PBS for 1 hour at 4°C, washed in PBS, dehydrated through an ethanol series and left overnight in 100% ethanol at 4°C. Samples were subjected to critical point drying (Leica EM CPD300) and gold-palladium sputter coating (Lnxor), then imaged on a Phenom Desktop SEM (Thermo Scientific). For phase contrast imaging, 800 μL of culture was centrifuged at 11000xg for 2 minutes and resuspended in 1 ml of fresh PBS. 5 μl of cell suspension was placed into a μslide chamber topped with a 2% low-melt agarose-PBS pad. Three biological replicates were imaged on a Leica DMi8 with a 100x phase contrast objective. Segmentation and quantification of cell morphology were done on approximately 10 non-overlapping images per sample using Nikon Nis Elements version 5.42.02 imaging software. A representative set of images from different genotypes was used to train the software using dynamic range adaptation and a final training loss of 0.00808. The resulting model was trained to ignore speckles, dividing cells, cells in clumps, touching neighboring cells, or image edges. Elongation is defined as the ratio between bacterial length and width.

## DATA AVAILABILITY

The atomic coordinates and structure factors have been deposited in the Protein Data Bank, www.rcsb.org, with the ID codes 9G58 and 9G59.

## ACKNOWLEDGEMENTS

Research was conducted at Center for High-Energy X-ray Sciences (CHEXS, MacCHESS), Cornell University Materials Research Science and Engineering Center (CCMR), and Advanced Photon Source (APS, NE-CAT beamlines). CHEXS and MacCHESS are supported by the NSF (DMR-2342336) and the NIH (P30-GM124166), respectively. CCMR is supported by the NSF (DMR-1719875). NE-CAT beamlines are supported by the NIH (P30-GM124165, S10-OD021527) at APS (DE-AC02-06CH11357). We thank John Grazul (CCMR, Cornell) and Roland Thünauer (Advanced Light and Fluorescence Microscopy Facility, CSSB) for providing training and technical support. This work was supported by the NIH (R01-AI114261 to F.H.Y. and R01-AI168017 to H.S.), and by BBSRC Institute Strategic Programme (BB/X010996/1 to J.G.M. and R.H.L.).

