## Supplemental Figures and Tables for "Secreted retropepsin-like enzymes are essential for stress tolerance and biofilm formation in *Pseudomonas aeruginosa*"

A

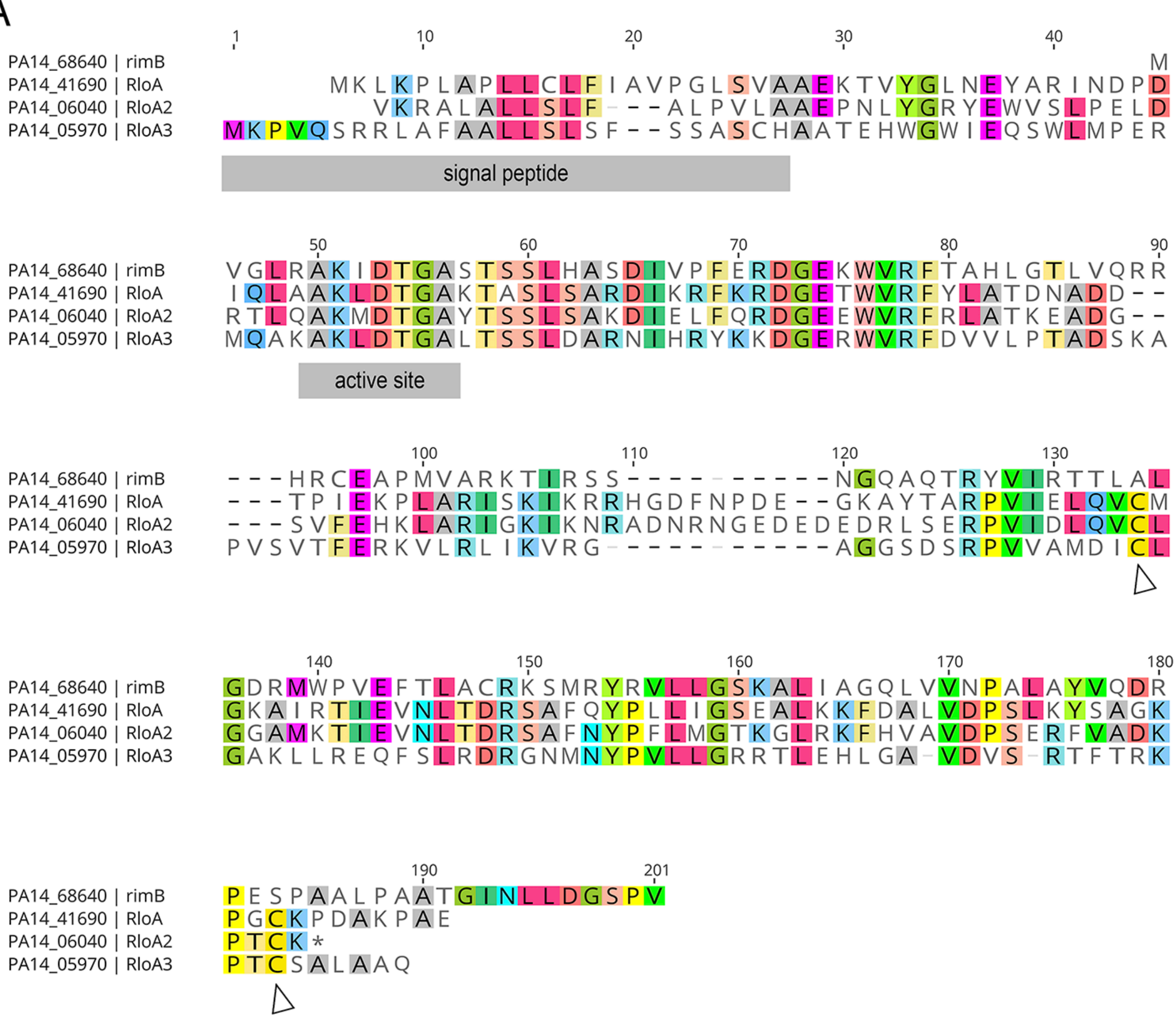


B

| | PA14_68640 \| rimB | PA14_41690 \| RloA | PA14_06040 \| RloA2 | PA14_05970 \| RloA3 |
|---|---|---|---|---|
| PA14_68640 \| rimB | | | | |
| PA14_41690 \| RloA | 30.99% | | | |
| PA14_06040 \| RloA2 | 31.88% | 50.87% | | |
| PA14_05970 \| RloA3 | 30.37% | 31.32% | 32.95% | |

**Figure S1.- Sequence conservation between RimB and RloA proteases**. **A.** Clustal Omega alignment highlighting the conserved signal peptide and conserved aspartic protease active site. Arrowheads point towards conserved disulfide-bond forming cysteine residues in RloA proteins. **B.** Protein sequence identity between pairwise combinations of RimB and RloA proteases.

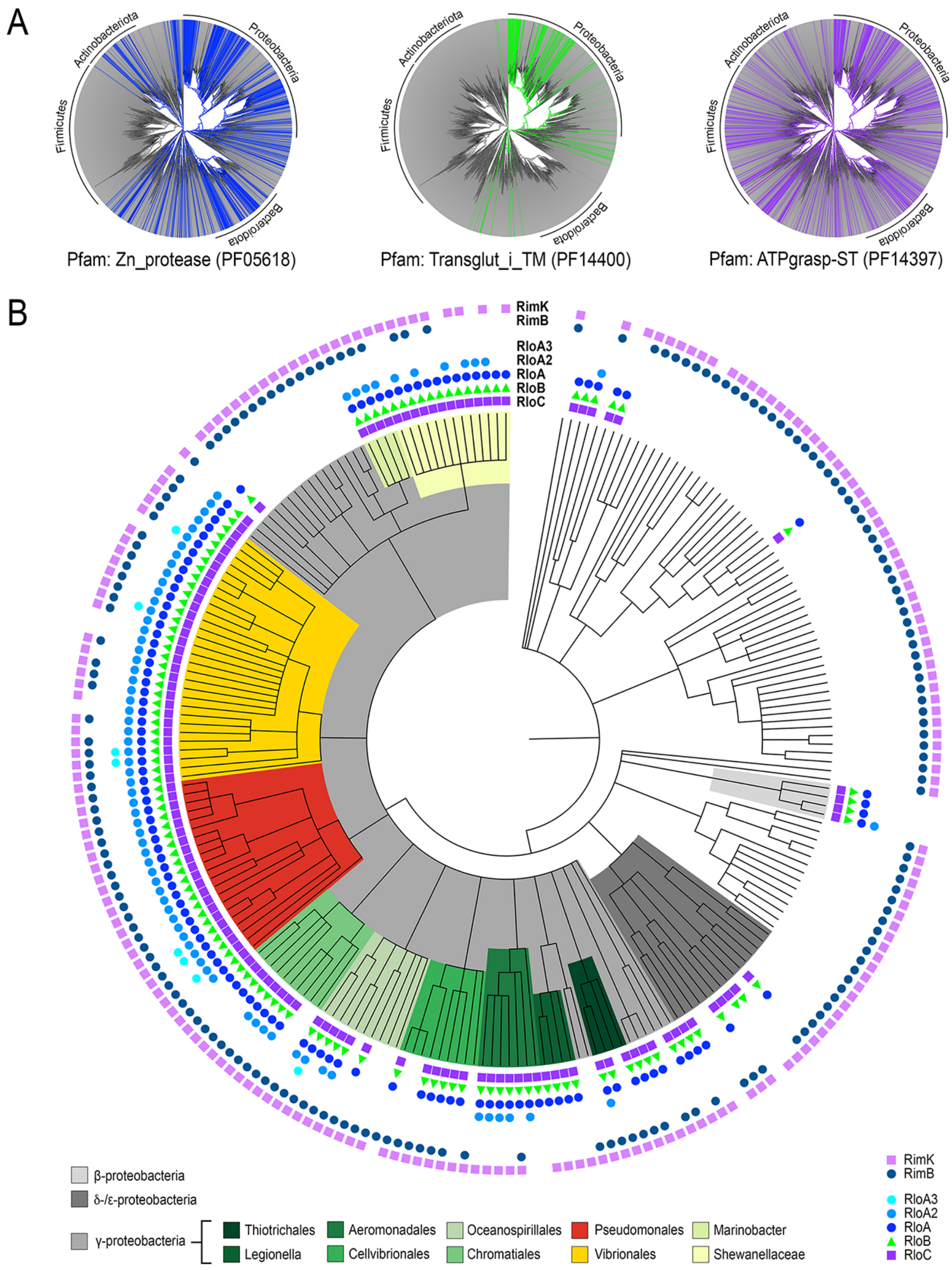


**Figure S2.- Phylogenetic analysis of RloA proteases and the *rlo* operon**. **A.** Phylogenetic conservation of Pfam groups for Zn_protease (PF05618, blue), Inactive transglutaminase fused to 7 transmembrane helices (PF1440, green) and ATP-grasp_ST (PF14397, purple) proteins is shown through AnnoTree plots (1). **B.** Plot showing the taxonomic distribution of RimB-RimK (outer circle) and Rlo proteins (inner circle). Plotted data was obtained by first surveying the EggNOG database (version 4.5.1) (2) with the RloA sequence as input. This retrieved a representative list of putative zinc protease sequences from bacteria (COG4067), which were then sorted according to their gene neighborhood into three groups: those adjacent to a *rimK* gene, those adjacent to *rloB* and *rloC* genes, and those with adjacent genes unrelated to the *rlo* or *rim loci*. The resulting gene clusters were then plotted using iTol (3). This analysis revealed that several organisms encode more than one RloA homologs containing N-terminal signal peptides (RloA2 and RloA3 in light blue circles). The fact that some organisms have more than one RloA homolog could either indicate emerging redundancy for RloA activity or further specialization within this enzyme family.

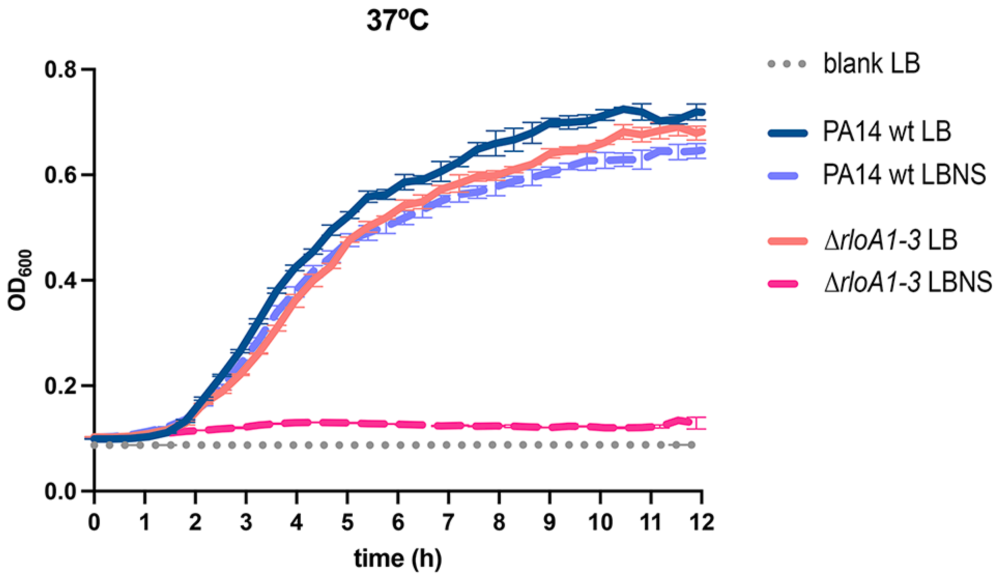


**Figure S3.- Growth curve PA14 wild type and ∆*rloA1-3* mutants.** Graph shows average $OD^{600}$ measurements of three experimental replicas for the bacterial strains and experimental conditions indicated in the legend. Error bars show standard deviation. LB - Lysogeny Broth, LBNS - LB without salts.

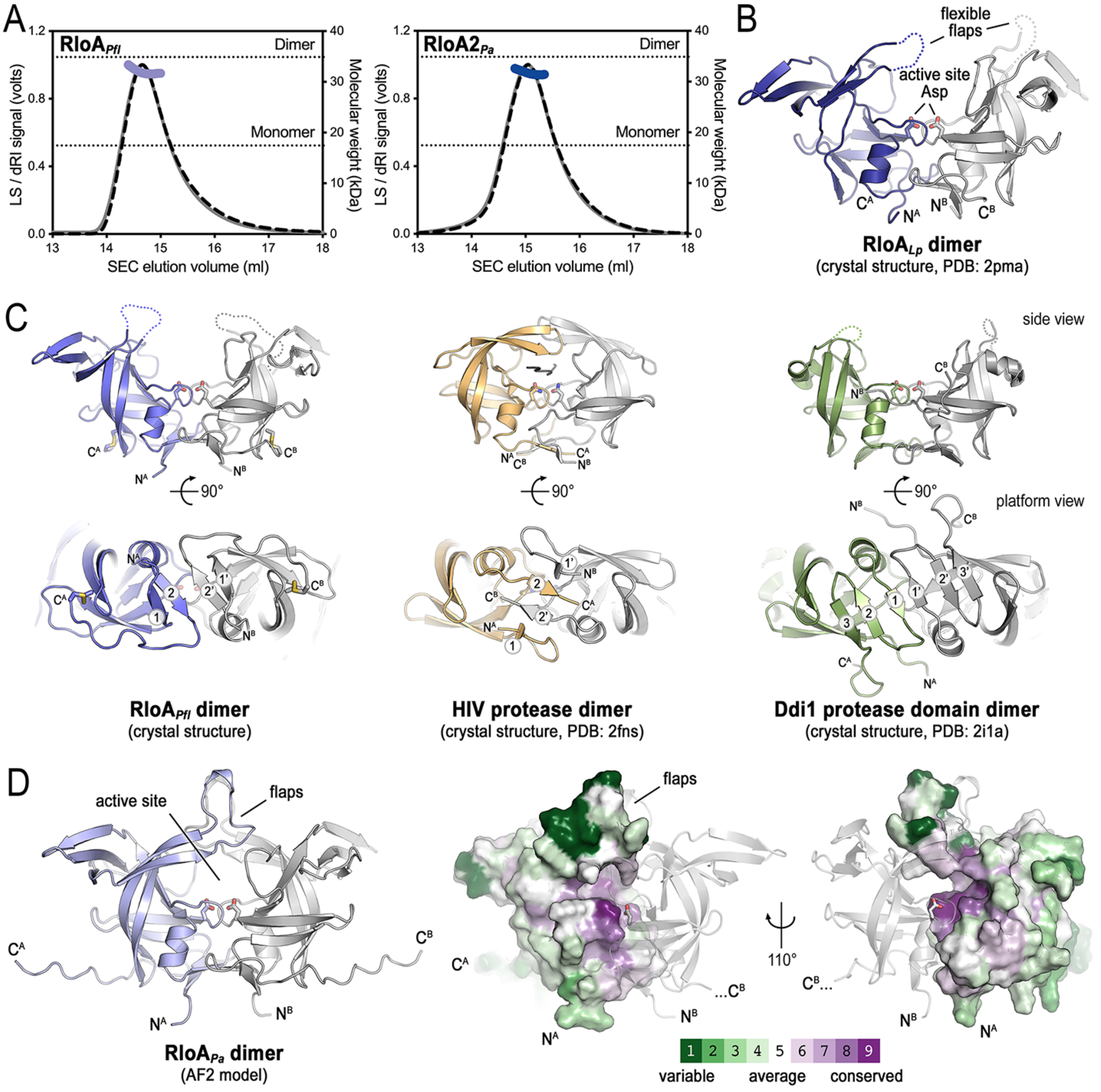


**Figure S4.- Structural features of RloA proteases**. **A.** SEC-MALS analysis of *P. fluorescens* RloA and *P. aeruginosa* RloA2. Normalized light scattering signal (grey line, right axis) and molar mass values (dashed line, left axis) were plotted against the SEC elution volume. Dotted lines show the theoretical molar mass ranges for monomeric and dimeric assemblies based on protein sequence. **B.** Crystal structure of a *Legionella pneumophila* RloA ortholog (PDB: 2pma). **C.** Crystal structures of *P. fluorescens* RloA (left), HIV protease (center, PDB: 2fns) and Ddi1 protease (right, PDB: 2i1a) dimers. Comparative side (top) and platform (bottom) views are shown. One of the protomers in each dimer has been colored in grey. **D.** ConSurf rendering of sequence conservation across a representative set of RloA proteases onto the AlphaFold2 model of *P. aeruginosa* RloA dimer. ConSurf scores are color coded as indicated. A ribbon view of the *P. aeruginosa* RloA AlphaFold2 model is shown to the left for reference.

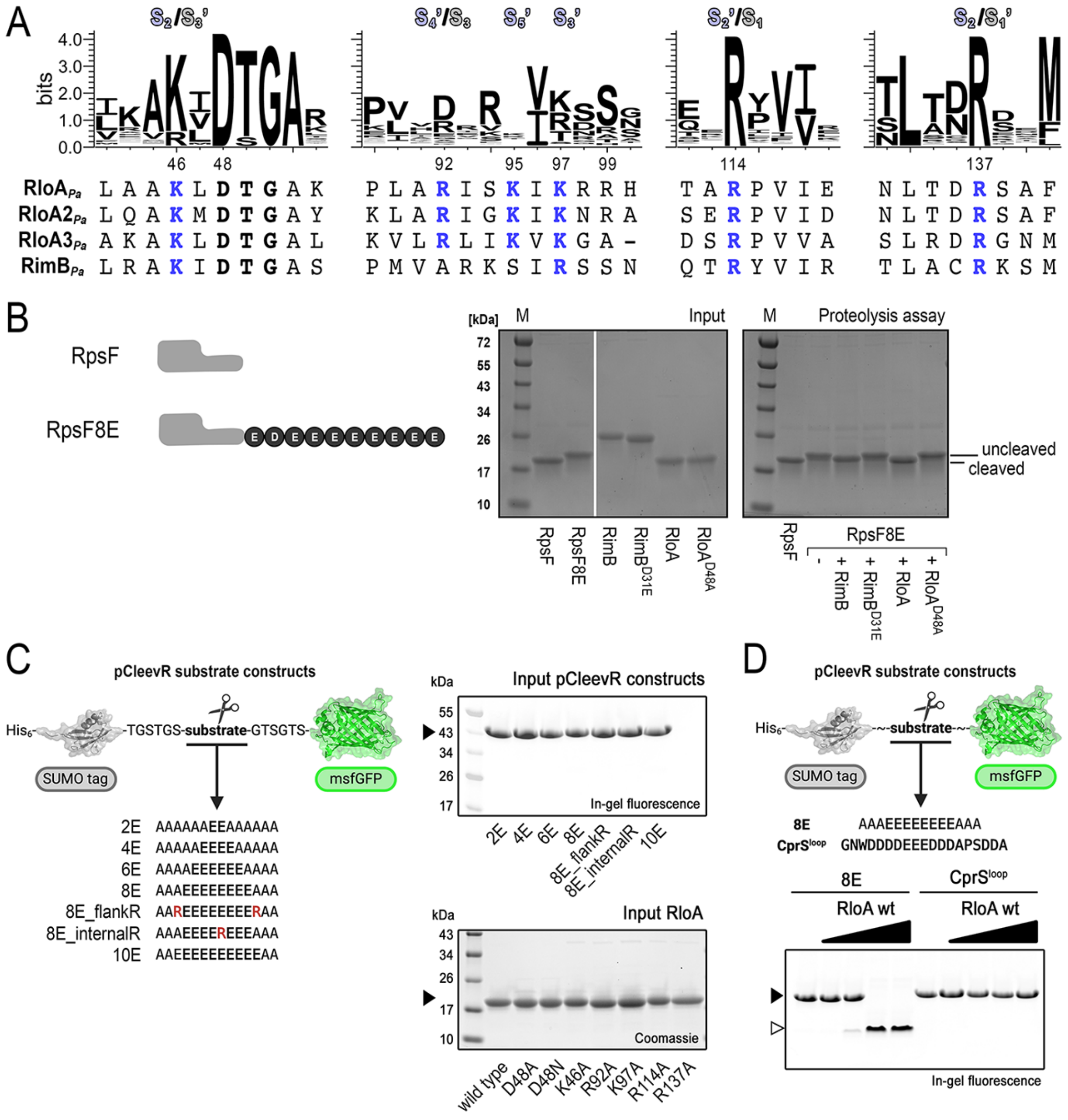

**Figure S5.- A.** Logos and alignments of RimB and RloA proteases highlighting the conserved active site (bold font) and residues involved in interactions with polyglutamate coordination in the AlphaFold2 model shown in Figure 3A (blue font). **B.** SDS-PAGE analysis showing the products of enzymatic reactions with the substrates schematically represented in the left panel and RimB or RloA proteins in their wild type or catalytically inactive mutant variants as indicated. Lines mark the positions of the cleaved and uncleaved substrates. **C.** Scheme illustrating the pCleevR polyglutamate substrates and input SDS-PAGE gels for experiments in Figure 3C-D. **D.** A sequence from CprS encoding a stretch of DE amino acids and their flanking residues was cloned into the pCleevR plasmid (top). Results from the incubation of RloA with the pCleevR-CprS$^{loop}$ reporter protein are shown at the bottom. The pCleevR-8E construct was used as control. Empty and filled arrowheads mark the positions of cleaved and uncleaved substrates, respectively.

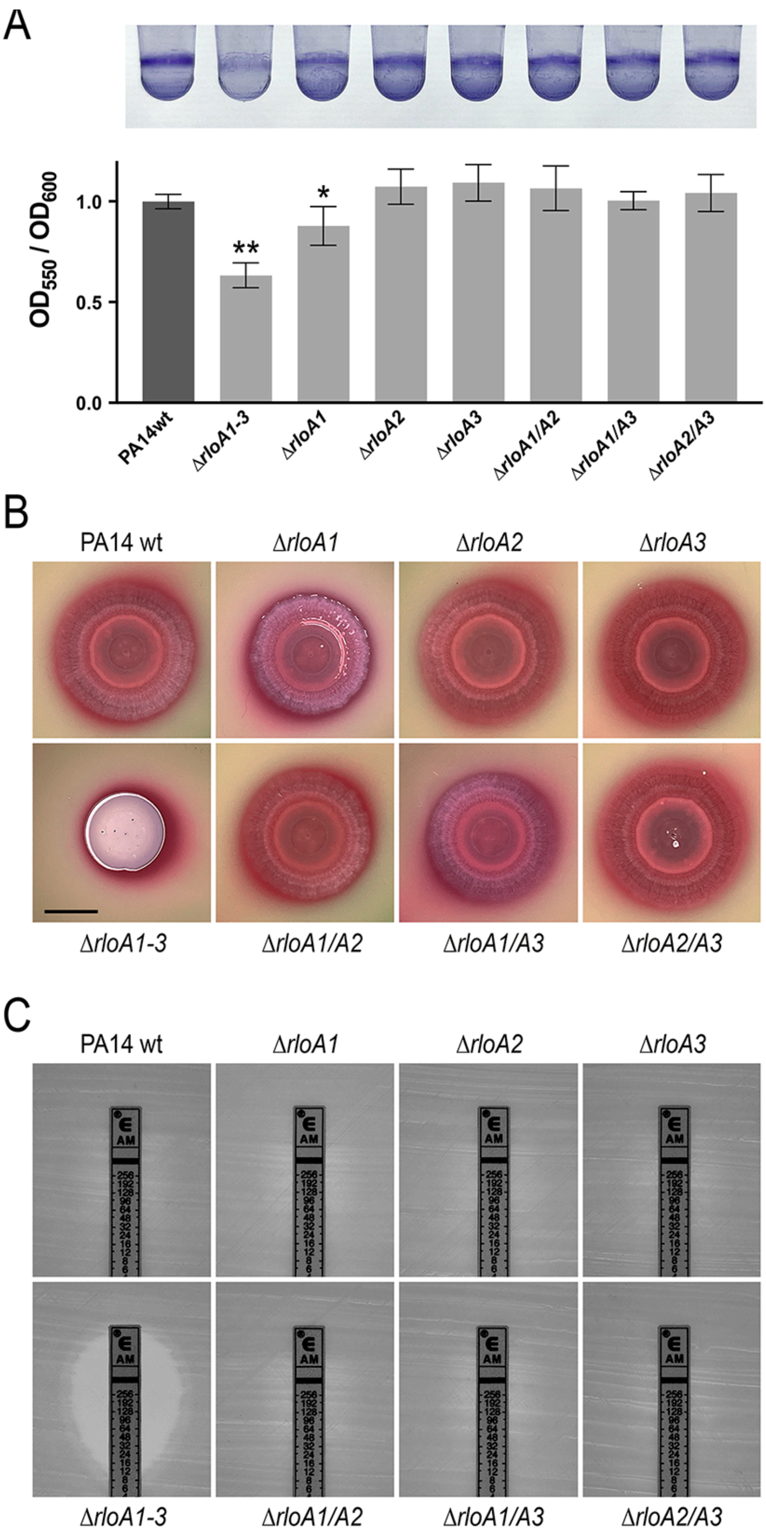


**Figure S6.- Phenotypic characterization of *rloA* mutant combinations. A.** Biofilm formation in PA14 wild type and combinations of RloA protease mutants was assessed through crystal violet staining of overnight cultures grown in 0.4% Arginine M63 media on PVC plates (top). Graph shows the normalized average intensity of crystal violet staining ($OD^{550}$) relative to the growth of the corresponding bacterial culture ($OD^{600}$), n=6. Error bars show standard deviation. Asterisks mark p values from paired two tailed t-tests against the PA14 wild type sample. *, p value <0.5. **, p value <0.0001. **B.** PA14 wild type and combinations of RloA protease mutant colonies grown in Congo Red 0.2% Glucose M63 agar plates. Scale bar represents 5 mm. **G.** Ampicillin ETEST strips tested on lawns of PA14 wild type and RloA protease mutant strains grown on Mueller-Hinton agar plates.

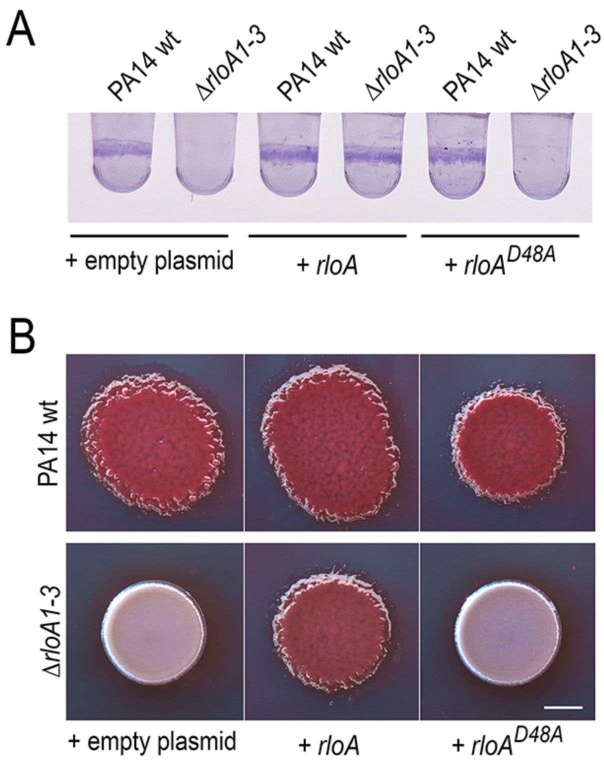


**Figure S7.- Biofilm growth in Δ*rloA1-3* mutants upon expression of plasmid-encoded wild type and catalytically inactive *rloA*. A.** Biofilm formation was assessed through crystal violet staining of overnight cultures grown on PVC plates in 0.2% arabinose M63 media with 25 μg/ml Gentamicin. **B.** Colonies of the indicated phenotypes were grown in Congo Red 0.4% Arginine M63 agar plates with 25 μg/ml Gentamicin. Scale bar represents 2 mm.

| | | |
|---|---|---|
| CELL WALL SYNTHESIS | | |
| AMPICILLIN | PA14 WT | RESISTANT ( >256.0 ) |
| AMPICILLIN | PA14 Δ*rloA1-3* | RESISTANT ( 16.0 ) |
| AUGMENTIN (AMOXICILLIN / CLAVULANIC ACID*) | PA14 WT | RESISTANT ( 128.0 ) |
| AUGMENTIN (AMOXICILLIN / CLAVULANIC ACID*) | PA14 Δ*rloA1-3* | RESISTANT ( 8.0 ) |
| PIPERACILLIN / TAZOBACT* | PA14 WT | SUSCEPTIBLE ( <=8.0 ) |
| PIPERACILLIN / TAZOBACT* | PA14 Δ*rloA1-3* | SUSCEPTIBLE ( <=8.0 ) |
| TICARCILLIN | PA14 WT | SUSCEPTIBLE ( 32.0 ) |
| TICARCILLIN | PA14 Δ*rloA1-3* | SUSCEPTIBLE ( 32.0 ) |
| CEFAZOLIN | PA14 WT | RESISTANT ( >32.0 ) |
| CEFAZOLIN | PA14 Δ*rloA1-3* | RESISTANT ( >32.0 ) |
| CEPHALEXIN | PA14 WT | RESISTANT ( >256.0 ) |
| CEPHALEXIN | PA14 Δ*rloA1-3* | RESISTANT ( 32.0 ) |
| CEFTAZIDIME | PA14 WT | SUSCEPTIBLE ( <=4.0 ) |
| CEFTAZIDIME | PA14 Δ*rloA1-3* | SUSCEPTIBLE ( <=4.0 ) |
| BACITRACIN | PA14 WT | RESISTANT ( >4.0 ) |
| BACITRACIN | PA14 Δ*rloA1-3* | RESISTANT ( >4.0 ) |
| PROTEIN SYNTHESIS - 50S RIBOSOMAL SUBUNIT | | |
| ERYTHROMYCIN | PA14 WT | RESISTANT ( >4.0 ) |
| ERYTHROMYCIN | PA14 Δ*rloA1-3* | RESISTANT ( >4.0 ) |
| CHLORAMPHENICOL | PA14 WT | RESISTANT ( >16.0 ) |
| CHLORAMPHENICOL | PA14 Δ*rloA1-3* | INTERMEDIATE ( 16.0 ) |
| PROTEIN SYNTHESIS - 30S RIBOSOMAL SUBUNIT | | |
| AMIKACIN | PA14 WT | SUSCEPTIBLE ( <=16.0 ) |
| AMIKACIN | PA14 Δ*rloA1-3* | SUSCEPTIBLE ( <=16.0 ) |
| GENTAMICIN | PA14 WT | SUSCEPTIBLE ( <=2.0 ) |
| GENTAMICIN | PA14 Δ*rloA1-3* | SUSCEPTIBLE ( <=2.0 ) |
| NEOMYCIN | PA14 WT | SUSCEPTIBLE ( <=4.0 ) |
| NEOMYCIN | PA14 Δ*rloA1-3* | SUSCEPTIBLE ( <=4.0 ) |
| TOBRAMYCIN | PA14 WT | SUSCEPTIBLE ( <=4.0 ) |
| TOBRAMYCIN | PA14 Δ*rloA1-3* | SUSCEPTIBLE ( <=4.0 ) |
| TETRACYCLINE | PA14 WT | RESISTANT ( 8.0 ) |
| TETRACYCLINE | PA14 Δ*rloA1-3* | RESISTANT ( 4.0 ) |
| DOXYCYCLINE | PA14 WT | RESISTANT ( >2.0 ) |
| DOXYCYCLINE | PA14 Δ*rloA1-3* | RESISTANT ( >2.0 ) |
| OXYTETRACYCLINE | PA14 WT | RESISTANT ( 4.0 ) |
| OXYTETRACYCLINE | PA14 Δ*rloA1-3* | RESISTANT ( 4.0 ) |
| NUCLEIC ACID SYNTHESIS | | |
| OFLOXACIN | PA14 WT | SUSCEPTIBLE ( 0.25 ) |
| OFLOXACIN | PA14 Δ*rloA1-3* | SUSCEPTIBLE ( 0.5 ) |
| ENROFLOXACIN | PA14 WT | SUSCEPTIBLE ( 0.5 ) |
| ENROFLOXACIN | PA14 Δ*rloA1-3* | SUSCEPTIBLE ( 0.5 ) |
| CIPROFLOXACIN | PA14 WT | SUSCEPTIBLE ( <=1.0 ) |
| CIPROFLOXACIN | PA14 Δ*rloA1-3* | SUSCEPTIBLE ( <=1.0 ) |
| TRIMETHOPRIM / SULFAMETHOXAZOLE | PA14 WT | RESISTANT ( 4.0 ) |
| TRIMETHOPRIM / SULFAMETHOXAZOLE | PA14 Δ*rloA1-3* | RESISTANT ( 4.0 ) |

**Figure S8.- Antibiotic susceptibility panel.** Table shows susceptibility of PA14 wild type and Δ*rloA1-3* mutants to a panel of antibiotics with different modes of action. β-lactamase inhibitors (asterisks) were included in some antibiotic test conditions.

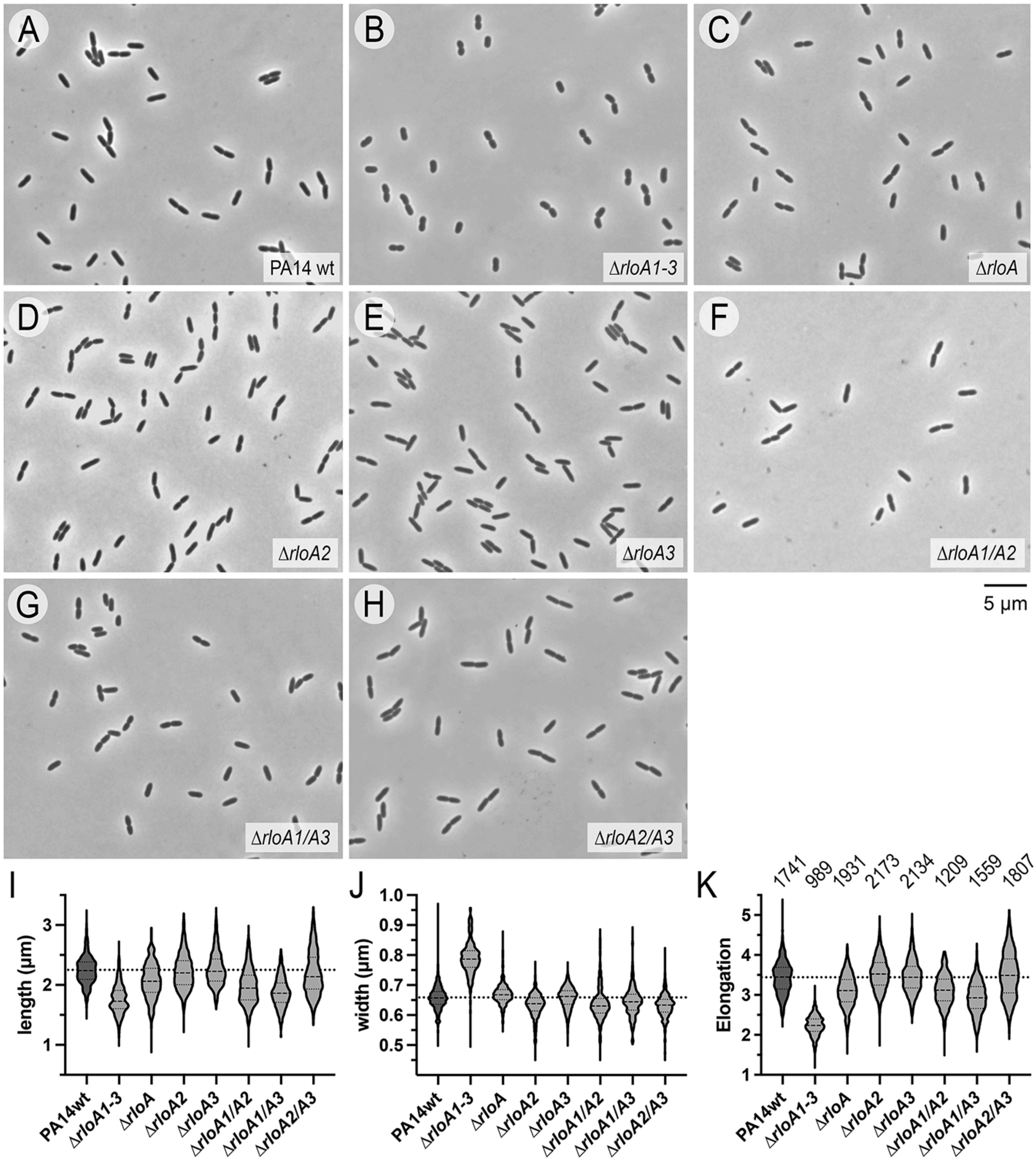


**Figure S9.- Cell morphology in RloA protease mutants. A-H.** Phase contrast images of PA14 wild type and RloA protease mutants of the indicated genotypes. Scale bar represents 5 µm. **I-K.** Violin plots showing quantification of cell length, width, and elongation. The number of samples analyzed for each strain is indicated on top of the graph in K. Quantification of measurements showed that individual Δ*rloA1* mutants, as well as double mutant combinations involving *rloA1* (Δ*rloA1/A2* and Δ*rloA1/A3* mutants) had a slight decrease in length. These results indicate that RloA1 might have a stronger dose-dependent requirement in comparison to the other RloA proteases.

**Table S1.** Crystalographic data collection and refinement statistics.*

| Data Collection | rloA_Pfl | rloA2_Pa |
|---|---|---|
| Wavelength (Å) | 0.9792 | 0.9792 |
| Resolution range (Å) | 45.0 - 2.28 (2.36 - 2.28) | 61.6 - 2.40 (2.47 - 2.40) |
| Space group | $P\ 4_1$ | $P\ 3_2\ 2\ 1$ |
| Unit cell dimensions *a*, *b*, *c* (Å) | 90.0 90.0 37.2 | 71.1 71.1 146.9 |
| Unit cell angles α, β, γ (°) | 90 90 90 | 90 90 120 |
| Total reflections | 148624 (13015) | 207392 (15389) |
| Unique reflections | 26253 (2402) | 32573 (2681) |
| Multiplicity | 5.7 (5.4) | 6.4 (5.7) |
| Completeness (%) | 98.91 (91.98) | 99.84 (98.48) |
| Mean I / σ(I) | 8.91 (1.42) | 16.45 (1.06) |
| $R_{merge}$ | 0.09684 (1.069) | 0.05016 (1.204) |
| CC1/2 | 0.994 (0.837) | 0.999 (0.498) |
| **Refinement** | | |
| Reflections used in refinement | 13726 (1262) | 17512 (1424) |
| Reflections used for R-free | 1363 (124) | 1750 (144) |
| $R_{work}$ / $R_{free}$ | 21.69 / 24.69 | 22.02 / 24.82 |
| Number of non-hydrogen atoms | 1978 | 2153 |
| macromolecules | 1951 | 2136 |
| solvent | 27 | 17 |
| RMS bonds (Å) | 0.003 | 0.002 |
| RMS angles (°) | 0.63 | 0.53 |
| Ramachandran favored (%) | 96.81 | 96.56 |
| Ramachandran allowed (%) | 3.19 | 3.44 |
| Ramachandran outliers (%) | 0 | 0 |
| Average B-factor | 71.48 | 77.65 |
| macromolecules | 71.69 | 77.72 |
| solvent | 56.1 | 68.19 |

*Values in parentheses are for highest-resolution shell.

**Table S2.** Bacterial strains used in this study.

| Strain | Genotype | Source/Reference | Additional information |
|---|---|---|---|
| *E.coli* T7 Express | | New England Biolabs | |
| *E. coli* NEB® 5α | | New England Biolabs | |
| *E. coli* S17.1 λpir$^{+}$ | | (4) | |
| *P. aeruginosa* UCBPP-PA14 | wild type (PA14wt) | (5) | TaxID 208963 |
| *P. aeruginosa* UCBPP-PA14 | Δ*rloA* | This study | Generated with pEX18-*rloA* on PA14wt strain |
| *P. aeruginosa* UCBPP-PA14 | Δ*rloA2* | This study | Generated with pEX18-*rloA2* on PA14wt strain |
| *P. aeruginosa* UCBPP-PA14 | Δ*rloA3* | This study | Generated with pEX18-*rloA3* on PA14wt strain |
| *P. aeruginosa* UCBPP-PA14 | Δ*rloA1/A2* | This study | Generated with pEX18-*rloA2* on Δ*rloA* strain |
| *P. aeruginosa* UCBPP-PA14 | Δ*rloA1/A3* | This study | Generated with pEX18-*rloA3* on Δ*rloA* strain |
| *P. aeruginosa* UCBPP-PA14 | Δ*rloA2/A3* | This study | Generated with pEX18-*rloA3* on Δ*rloA2* strain |
| *P. aeruginosa* UCBPP-PA14 | Δ*rloA1-3* | This study | Generated with pEX18-*rloA3* on Δ*rloA1/A2* strain |
| *P. aeruginosa* UCBPP-PA14 | Δ*rloB* | This study | Generated with pEX18-*rloB* on PA14wt strain |
| *P. aeruginosa* UCBPP-PA14 | Δ*rloC* | This study | Generated with pEX18-*rloC* on PA14wt strain |
| *P. aeruginosa* UCBPP-PA14 | Δ*rloABC* | This study | Generated with pEX18-*rloABC* on PA14wt strain |
| *P. aeruginosa* UCBPP-PA14 | Δ*rloA1-3;* Δ*rloB* | This study | Generated with pEX18-*rloAB* on Δ*rloA1/A2* strain |
| *P. aeruginosa* UCBPP-PA14 | Δ*rloA1-3;* Δ*rloC* | This study | Generated with pEX18-*rloC* on Δ*rloA1-3* strain |
| *P. aeruginosa* UCBPP-PA14 | Δ*rloA1-3;* Δ*rloBC* | This study | Generated with pEX18-*rloABC* on Δ*rloA1/A2* strain |
| *P. aeruginosa* UCBPP-PA14 | Δ*rimB* | This study | Generated with pEX18-*rimB* on PA14wt strain |
| *P. aeruginosa* UCBPP-PA14 | Δ*rloA1-3;* Δ*rimB* | This study | Generated with pEX18-*rimB* on Δ*rloA1-3* strain |
| *P. aeruginosa* UCBPP-PA14 | Δ*rloA1/A2;* Δ*rimB* | This study | Generated with pEX18-*rimB* on Δ*rloA1/A2* strain |
| *P. aeruginosa* UCBPP-PA14 | Δ*rloA1/A3;* Δ*rimB* | This study | Generated with pEX18-*rimB* on Δ*rloA1/A3* strain |
| *P. aeruginosa* UCBPP-PA14 | Δ*rloA2/A3;* Δ*rimB* | This study | Generated with pEX18-*rimB* on Δ*rloA2/A3* strain |

**Table S3.** Plasmids used in this study.

| Plasmid | Description | Source |
|---|---|---|
| $His_6$-SUMO-pET28 | Fusion proteins cloned into this pET28-derived vector via BamHI-NotI render the protein of interest linked to an N-terminal His6-tagged small ubiquitin-like modifier (SUMO) cleavable by recombinant Ulp-1 protease. Antibiotic selection in 50 µg/ml Kanamycin | This study |
| $His_6$-SUMO-pET28-PA1768(23-179) | $RloA_{Pa}$ (aa 23-179 from PA1768) was PCR amplified and cloned via BamHI-NotI into His6-SUMO-pET28 | This study |
| $His_6$-SUMO-pET28-PA1768(23-179) - D48A | Site directed mutagenesis was performed on $His_6$-SUMO-pET28-PA1768(23-179) | This study |
| $His_6$-SUMO-pET28-PA1768(23-179) - D48N | Site directed mutagenesis was performed on $His_6$-SUMO-pET28-PA1768(23-179) | This study |
| $His_6$-SUMO-pET28-PA1768(23-179) - K46A | Site directed mutagenesis was performed on $His_6$-SUMO-pET28-PA1768(23-179) | This study |
| $His_6$-SUMO-pET28-PA1768(23-179) - R92A | Site directed mutagenesis was performed on $His_6$-SUMO-pET28-PA1768(23-179) | This study |
| $His_6$-SUMO-pET28-PA1768(23-179) - K97A | Site directed mutagenesis was performed on $His_6$-SUMO-pET28-PA1768(23-179) | This study |
| $His_6$-SUMO-pET28-PA1768(23-179) - R114A | Site directed mutagenesis was performed on $His_6$-SUMO-pET28-PA1768(23-179) | This study |
| $His_6$-SUMO-pET28-PA1768(23-179) - R137A | Site directed mutagenesis was performed on $His_6$-SUMO-pET28-PA1768(23-179) | This study |
| $His_6$-SUMO-pET28-PA0462(18-169) | $RloA2_{Pa}$ (aa 18-169 from PA0462)was PCR amplified and cloned via BamHI-NotI into His6-SUMO-pET28 | This study |
| $His_6$-SUMO-pET28-Pfl01_1762(20-178) | $RloA_{Pf}$ (aa 20-178 from Pfl01_1762) was PCR amplified and cloned via BamHI-NotI into His6-SUMO-pET28 | This study |
| pETM11-RimB | | (6) |
| pETM11-$RimB^{D31E}$ | | (7) |
| pETM11-RpsF | | (6) |
| pETM11-RpsF8E | | (7) |
| pCleevR | Antibiotic selection in 50 µg/ml Kanamycin | (10) |
| pCleevR-2E | 2E gene fragment was cloned via BamHI-KpnI into pCleevR | This study |
| pCleevR-4E | 4E gene fragment was cloned via BamHI-KpnI into pCleevR | This study |
| pCleevR-6E | 6E gene fragment was cloned via BamHI-KpnI into pCleevR | This study |
| pCleevR-8E | 8E gene fragment was cloned via BamHI-KpnI into pCleevR | This study |
| pCleevR-10E | 10E gene fragment was cloned via BamHI-KpnI into pCleevR | This study |
| pCleevR-8E_flankR | 8E_flankR gene fragment was cloned via BamHI-KpnI into pCleevR | This study |
| pCleevR-8E_internalR | 8E_internalR gene fragment was cloned via BamHI-KpnI into pCleevR | This study |
| pCleevR-$CprS^{loop}$ | $CprS^{loop}$ gene fragment was cloned via BamHI-KpnI into pCleevR | This study |

**Table S3 (cont.)** Plasmids used in this study.

| Plasmid | Description | Source |
|---|---|---|
| pJN105 | Antibiotic selection in 15 µg/ml Gentamicin (*E.coli*) or 25-50 µg/ml Gentamicin (*Pseudomonas aeruginosa* PA14) | (8) |
| pJHA | Antibiotic selection in 15 µg/ml Gentamicin (*E.coli*) or 25-50 µg/ml Gentamicin (*Pseudomonas aeruginosa* PA14) | (9) |
| pJHA-*rloA* | *rloA* (PA14_41690) was PCR amplified and cloned via NdeI-EcoRI into pJHA | This study |
| pJHA-*rloA*$^{D48A}$ | Site directed mutagenesis of pJHA-*rloA* | This study |
| pJHA-*rloA*$^{K46A}$ | Site directed mutagenesis of pJHA-*rloA* | This study |
| pJHA-*rloA*$^{R92A}$ | Site directed mutagenesis of pJHA-*rloA* | This study |
| pJHA-*rloA*$^{K97A}$ | Site directed mutagenesis of pJHA-*rloA* | This study |
| pJHA-*rloA*$^{R114A}$ | Site directed mutagenesis of pJHA-*rloA* | This study |
| pJHA-*rloA*$^{R137A}$ | Site directed mutagenesis of pJHA-*rloA* | This study |
| pJHA-*rloA2* | *rloA2* (aa 156-234 from PA14_06040)[1] was PCR amplified and cloned via NdeI-EcoRI into pJHA | This study |
| pJHA-*rloA3* | *rloA3* (PA14_05970) was PCR amplified and cloned via NdeI-EcoRI into pJHA | This study |
| pJStrep | Strep-NheI-XbaI gene fragment was cloned via NheI-XbaI into pJN105. Antibiotic selection in 15 µg/ml Gentamicin (*E.coli*) or 25-50 µg/ml Gentamicin (*Pseudomonas aeruginosa* PA14) | This study |
| pJGCStrepII | msfGFP_CstrepII gene fragment was cloned via EcoRI-XbaI into pJN105. The EcoRI site at the polylinker was destroyed during cloning. pJGCStrepII can be used to generate C terminal Strep-tagged constructs (via EcoRI-HindIII) or proteins tagged with msfGFP at the N terminus and with Strep-tagII at the C terminus (via HindIII). Antibiotic selection in 15 µg/ml Gentamicin (*E.coli*) or 25-50 µg/ml Gentamicin (*Pseudomonas aeruginosa* PA14) | This study |
| pJStrep-*rloA* | *r*loA_CstrepII gene fragment was cloned via EcoRI-XbaI into pJN105 | This study |
| pJStrep-*rloA*$^{D48A}$ | Site directed mutagenesis of pJStrep-*rloA* | This study |
| pEX18Gm | Antibiotic selection in 15 µg/ml Gentamicin (*E.coli*) or 50 µg/ml Gentamicin (*Pseudomonas aeruginosa* PA14) | (4) |
| pEX18-*rloA* | *rloA* deletion allele was amplified through overlap extension PCR, then cloned via EcoRI-BamHI into pEX18-Gm | This study |
| pEX18-*rloA2* | *rloA2* deletion allele was amplified through overlap extension PCR, then cloned via EcoRI-BamHI into pEX18-Gm | This study |
| pEX18-*rloA3* | *rloA3* deletion allele was amplified through overlap extension PCR, then cloned via EcoRI-BamHI into pEX18-Gm | This study |
| pEX18-*rloB* | *rloB* deletion allele was amplified through overlap extension PCR, then cloned via EcoRI-BamHI into pEX18-Gm | This study |
| pEX18-*rloC* | *rloC* deletion allele was amplified through overlap extension PCR, then cloned via EcoRI-BamHI into pEX18-Gm | This study |
| pEX18-*rloABC* | *rlo operon* deletion allele was amplified through overlap extension PCR, then cloned via EcoRI-BamHI into pEX18-Gm | This study |
| pEX18-*rloAB* | *rloAB* deletion allele was amplified through overlap extension PCR, then cloned via EcoRI-BamHI into pEX18-Gm | This study |
| pEX18-*rimB* | *rimB* deletion allele was amplified through overlap extension PCR, then cloned via EcoRI-BamHI into pEX18-Gm | This study |

[1] Note that PA14_06040 is annotated in Pseudomonas.com as a longer transcript that overlaps with gene PA14_06030 and is predicted to encode a protein without a signal peptide. However, orthologous genes in other *Pseudomonas* species (i.e. *P. aeruginosa* PA7 and *P. fluorescens* SBW25 at Pseudomonas.com or *P. aeruginosa* PAO1 at KEGG) are annotated as shorter transcripts encoding a signal peptide. We have considered PA14 RloA2 to encode an equivalent short protein of 169aa with a signal peptide. Here we refer to RloA2 as containing amino acids 156 to 234 of the annotated version of PA14_06040 available at Pseudomonas.com at the time this manuscript was written.

**Table S4.** Primers used in this study.

| Name | Sequence | Purpose |
|---|---|---|
| PA1768-RloA-23BamHI-F | GAATTCGGATCCGCCGAAAAGACGGTCTACGGC | Insert for $His_6$-SUMO-pET28-PA1768(23-179) |
| PA1768-RloA-179stopNotI-R | CTCGAGGCGGCCGCTTACTCAGCAGGTTTTGCATCAGG | Insert for $His_6$-SUMO-pET28-PA1768(23-179) |
| PA0462s-RloA2-18-SUMO-F | ACAGATTGGTGGATCCGCCGAACCCAACCTCTACG | Insert for $His_6$-SUMO-pET28-PA0462(18-169) |
| PA0462s-RloA2-169-SUMO-R | TGCTCGAGTGCGGCCGCTTACTTGCAGGTGGGCTTGTC | Insert for $His_6$-SUMO-pET28-PA0462(18-169) |
| Pfl1762-20BamHI-F | CATATGGGATCCGCGGCGGGGGAAAAGACCGTGTACG | Insert for $His_6$-SUMO-pET28-Pfl01_1762(20-178) |
| Pfl1762-178stopNotI-R | TCTAGAGCGGCCGCTTACTCGGCGGTATGAGCGGCGATGG | Insert for $His_6$-SUMO-pET28-Pfl01_1762(20-178) |
| PA14_41690-RloA-1NdeI-F | GGATCCCATATGAAACTCAAGCCCTTAGCTCC | Insert for pJHA-*rloA* |
| PA14_41690-RloA-EcoRI-R | CGTCAGAATTCCTCAGCAGGTTTTGCATCAGGTTTGC | Insert for pJHA-*rloA* |
| rloA_D48A_s | CGCCAAGCTCGCCACCGGCGCCA | *rloA* D48A mutagenesis |
| rloA_D48A_as | TGGCGCCGGTGGCGAGCTTGGCG | *rloA* D48A mutagenesis |
| rloA_D48N_s | CCGCCAAGCTCAACACCGGCGCC | *rloA* D48N mutagenesis |
| rloA_D48N_as | GGCGCCGGTGTTGAGCTTGGCGG | *rloA* D48N mutagenesis |
| RloA_K46A_s | GCCGGTGTCGAGCGCGGCGGCCAGCTGG | *rloA* K46A mutagenesis |
| RloA_K46A_as | CCAGCTGGCCGCCGCGCTCGACACCGGC | *rloA* K46A mutagenesis |
| RloA_R92A_s | TTGATCTTGCTGATCGCCGCCAGGGGCTTCTC | *rloA* R92A mutagenesis |
| RloA_R92A_as | GAGAAGCCCCTGGCGGCGATCAGCAAGATCAA | *rloA* R92A mutagenesis |
| RloA_K97A_s | CCGTGGCGGCGCGCGATCTTGCTGATCCGCG | *rloA* K97A mutagenesis |
| RloA_K97A_as | CGCGGATCAGCAAGATCGCGCGCCGCCACGG | *rloA* K97A mutagenesis |
| RloA_R114A_s | CTCGATCACCGGAGCGGCGGTATAGGCC | *rloA* R114A mutagenesis |
| RloA_R114A_as | GGCCTATACCGCCGCTCCGGTGATCGAG | *rloA* R114A mutagenesis |
| RloA_R137A_s | GGGTATTGGAATGCACTTGCGTCGGTCAGGTTCACTTC | *rloA* R137A mutagenesis |
| RloA_R137A_as | GAAGTGAACCTGACCGACGCAAGTGCATTCCAATACCC | *rloA* R137A mutagenesis |
| PA14_06040-RloA2-F | AAGGAGATATACATATGAAGCGTGCCCTTGCCTTG | Insert for pJHA-*rloA2* |
| PA14_06040-RloA2-EcoRI-R | CGTCAGAATTCCTTGCAGGTGGGCTTGTCGGCGACG | Insert for pJHA-*rloA2* |
| PA14_05970-RloA3-F | AAGGAGATATACATATGAAGCCTGTGCAATCCCGTCG | Insert for pJHA-*rloA3* |
| PA14_05970-RloA3-EcoRI-R | CGTCAGAATTCCTGAGCGGCGAGCGCGCTGCAGG | Insert for pJHA-*rloA3* |
| PA14_41690-RloA-UF | CCATGATTACGAATTCCGAGGTACGCGAGGTCGA | Insert for pEX18-*rloA* |
| PA14_41690-RloA-UR | GATGCAGGTTGAGAGAGCGCATGTCGTAAGTCCTGGTTGGTGTCGCGTTCGGC | Insert for pEX18-*rloA* |
| PA14_41690-RloA-DF | ACGACATGCGCTCTCTCAACCTGCATC | Insert for pEX18-*rloA* |
| PA14_41690-RloA-DR | CGACTCTAGAGGATCCGGCTGTCGCGGAAGATCG | Insert for pEX18-*rloA* |
| rloA-seqF | ATCGACCCCGATCGCCTCA | Confirm allele deletion |
| rloA-seqR | CTGACGAAGGTCTCGACGTC | Confirm allele deletion |
| PA14_06040-RloA2-UF | CGACTCTAGAGGATCCGGAAGGGCACCCAGATCAG | Insert for pEX18-*rloA2* |
| PA14_06040-RloA2-UR | TTACTTGCAGGTGGGCTTGTCCAAGGCAAGGGCACGCTTCAC | Insert for pEX18-*rloA2* |
| PA14_06040-RloA2-DF | GACAAGCCCACCTGCAAGTAA | Insert for pEX18-*rloA2* |
| PA14_06040-RloA2-DR | CCATGATTACGAATTCGATCCGCACGCGCTCCTC | Insert for pEX18-*rloA2* |

**Table S4 (cont.)** Primers used in this study.

| Name | Sequence | Purpose |
|---|---|---|
| rloA2-seqF | CTTGGTGATCTCCAGGTC | Confirm allele deletion |
| rloA2-seqR | CTGCTCGCGGCTGAATAC | Confirm allele deletion |
| PA14_05970-RloA3-UF | CGACTCTAGAGGATCCGTCGTAGAGTGCTGACACTG | Insert for pEX18-*rloA3* |
| PA14_05970-RloA3-UR | CTACTGAGCGGCGAGCGCGCTACGGGATTGCACAGGCTTCAT | Insert for pEX18-*rloA3* |
| PA14_05970-RloA3-DF | AGCGCGCTCGCCGCTCAGTAG | Insert for pEX18-*rloA3* |
| PA14_05970-RloA3-DR | CCATGATTACGAATTCCTGATCCTCGGCTGCCTG | Insert for pEX18-*rloA3* |
| rloA3-seqF | GCAGCGACAGGCAATGTA | Confirm allele deletion |
| rloA3-seqR | ATGACCATGATCCCGCTG | Confirm allele deletion |
| PA14_41710-RloB-UF | CCATGATTACGAATTCCCTTAGCTCCCCTGCTCTGC | Insert for pEX18-*rloB* |
| PA14_41710-RloB-UR | CGCGGTCAGGTCTTTCCAGGTCTTCCAGATCGACATGGTTCAGTCCTTGAGCAGGGCCTT ATGCAGGTTGAGAGAGCGCATGTCGTTTACTCAGCAGGTTTTGCATCAGG | Insert for pEX18-*rloB* |
| PA14_41710-RloB-DF | CCATGTCGATCTGGAAGACCTGGAAAGACCTGACCGCG | Insert for pEX18-*rloB* |
| PA14_41710-RloB-DR | CGACTCTAGAGGATCCTCGGCAGGCGCAGCATGG | Insert for pEX18-*rloB* |
| rloB-seqF | GCGTGATAGAATGCGGCC | Confirm allele deletion |
| rloB-seqR | GACAACTCGTAGCAACCGG | Confirm allele deletion |
| PA14_41730-RloC-UF | CCATGATTACGAATTCGCGCTCCTATCTCGAGCA | Insert for pEX18-*rloC* |
| PA14_41730-RloC-UR | GCCGGGCTTCTTCATACCGGAAGCACGGGTTCAGTCCTTGAGCAGGGCCTTGAAGCGG | Insert for pEX18-*rloC* |
| PA14_41730-RloC-DF | CGTGCTTCCGGTATGAAGAAGCCCGGC | Insert for pEX18-*rloC* |
| PA14_41730-RloC-DR | CGACTCTAGAGGATCCCCCAGGGTGTCTACGGCC | Insert for pEX18-*rloC* |
| rloC-seqF | CGGTTCTGATCGCCCTGG | Confirm allele deletion |
| rloC-seqR | CCGCGCTACGAGGAGTTG | Confirm allele deletion |
| PA14_41690to41730-rloABC-UF | CGACTCTAGAGGATCCGGAACAGGAGCTTTCGGCC | Insert for pEX18-*rloABC* |
| PA14_41690to41730-rloABC-UR | TCAGCCCTGGTGCTGCTGGACAGCTAAGGGCTTGAGTTTCAT | Insert for pEX18-*rloABC* |
| PA14_41690to41730-rloABC-DF | GTCCAGCAGCACCAGGGCTGA | Insert for pEX18-*rloABC* |
| PA14_41690to41730-rloABC-DR | CCATGATTACGAATTCCCCTACTACACCTGGGTGG | Insert for pEX18-*rloABC* |
| PA14_41690+41710-rloAB-UR | TCAGTCCTTGAGCAGGGCCTTAGCTAAGGGCTTGAGTTTCAT | Insert for pEX18-*rloAB* |
| PA14_41690+41710-rloAB-DF | AAGGCCCTGCTCAAGGACTGA | Insert for pEX18-*rloAB* |
| PA14_41690+41710-rloAB-DR | CCATGATTACGAATTCGGGCGTTGAGTTCGAGGATCA | Insert for pEX18-*rloAB* |
| rloAB-seqF | GCTTCGTCAAACTGACGAG | Confirm allele deletion |
| rloAB-seqF | GGTGCGTTGAAATCCGAG | Confirm allele deletion |
| PA14_68640-RimB-BamHI-UF | CAGGTCGACTCTAGAGGATCCCGCAACTGGTGGATCTGCC | Insert for pEX18-*rimB* |
| PA14_68640-RimB-UR | AAACCGGACTCCCGACCATGCCCAGTTC | Insert for pEX18-*rimB* |
| PA14_68640-RimB-DF | CATGGTCGGGAGTCCGGTTTGAGCCATTCG | Insert for pEX18-*rimB* |
| PA14_68640-RimB-EcoRI-DR | TATGACCATGATTACGAATTCTCCAGCAGCTTGATCACC | Insert for pEX18-*rimB* |
| rimB-seqF | CTGGCCGTGCTGTTCATG | Confirm allele deletion |
| rimB-seqR | CCTGGACCATGATGTTGTGC | Confirm allele deletion |

**Table S5.** Gene synthesized DNA fragments used in this study.

| Name | Sequence | Purpose |
|---|---|---|
| 2E | **TGGAAGCACGGGATCC**GCAGCTGCTGCGGCAGCCGAAGAGGCTGCA<br>GCTGCTGCAGCG**GGTACCAGCGGAACTTCTAG** | Insert for pCleevR-2E |
| 4E | **TGGAAGCACGGGATCC**GCAGCTGCTGCGGCAGAGGAAGAGGAAGCA<br>GCTGCTGCAGCG**GGTACCAGCGGAACTTCTAG** | Insert for pCleevR-4E |
| 6E | **TGGAAGCACGGGATCC**GCAGCTGCTGCGGAAGAGGAAGAGGAAGAA<br>GCTGCTGCAGCG**GGTACCAGCGGAACTTCTAG** | Insert for pCleevR-6E |
| 8E | **TGGAAGCACGGGATCC**GCAGCTGCTGAGGAAGAGGAAGAGGAAGAA<br>GAGGCTGCAGCG**GGTACCAGCGGAACTTCTAG** | Insert for pCleevR-8E |
| 10E | **TGGAAGCACGGGATCC**GCAGCTGAAGAGGAAGAGGAAGAGGAAGA<br>AGAGGAGGCAGCG**GGTACCAGCGGAACTTCTAG** | Insert for pCleevR-10E |
| 8E_flankR | **TGGAAGCACGGGATCC**GCAGCTAGAGAAGAAGAGGAAGAGGAAGA<br>GGAAAGAGCTGCA**GGTACCAGCGGAACTTCTAG** | Insert for pCleevR-8E_flankR |
| 8E_internalR | **TGGAAGCACGGGATCC**GCAGCTGCTGAGGAAGAGGAAAGAGAAGAA<br>GAGGCTGCAGCG**GGTACCAGCGGAACTTCTAG** | Insert for pCleevR-8E_internalR |
| CprS$^{loop}$ | **TGGAAGCACGGGATCC**GGTAACTGGGACGATGATGACGAAGAGGAA<br>GATGACGATGCGCCGTCTGACGATGCG**GGTACCAGCGGAACTTCTAG** | Insert for pCleevR-CprS$^{loop}$ |
| Strep-NheI-XbaI | **GCTAGC**CATATGCCCGGGGAGCTCGAATTCAAGCTTGAGCTCCCT**TGG**<br>**AGCCACCCGCAGTTCGAGAAG**TAA**TCTAGA** | Insert for pJStrep |
| msfGFP_CstrepII | **TTGGGCTAGCGAATT**GAATTC**ATG**AGCAAAGGTGAAGAACTGTTTACC<br>GGCGTTGTGCCGATTCTGGTGGAACTGGATGGTGATGTGAATGGCCA<br>TAAATTTAGCGTTCGTGGCGAAGGCGAAGGTGATGCGACCAACGGTA<br>AACTGACCCTGAAATTTATTTGCACCACCGGTAAACTGCCGGTTCCGTG<br>GCCGACCCTGGTGACCACCCTGACCTATGGCGTTCAGTGCTTTAGCCG<br>CTATCCGGATCATATGAAACGCCATGATTTCTTTAAAAGCGCGATGCCG<br>GAAGGCTATGTGCAGGAACGTACCATTAGCTTCAAAGATGATGGCACC<br>TATAAAACCCGTGCGGAAGTTAAATTTGAAGGCGATACCCTGGTGAAC<br>CGCATTGAACTGAAAGGTATTGATTTTAAAGAAGATGGCAACATTCTG<br>GGTCATAAACTGGAATATAATTTCAACAGCCATAATGTGTATATTACCG<br>CCGATAAACAGAAAAATGGCATCAAAGCGAACTTTAAAATCCGTCACA<br>ACGTGGAAGATGGTAGCGTGCAGCTGGCGGATCATTATCAGCAGAAT<br>ACCCCGATTGGTGATGGCCCGGTGCTGCTGCCGGATAATCATTATCTG<br>AGCACCCAGAGCAAACTGAGCAAAGATCCGAATGAAAAACGTGATCA<br>TATGGTGCTGCTGGAATTTGTTACCGCCGCGGGCATTACCCACGGTAT<br>GGATGAACTGTATAAAGGCAGCAAGCTTTCTGCGTGGTCTCACCCGCA<br>GTTCGAAAAA**TAA**TCTAGA**GCGGCCGCCA** | Insert for pJGCStrepII |
| rloA_CstrepII | **TTGGGCTAGCGAATTCATG**AAACTCAAGCCCTTAGCTCCCCTGCTCTGC<br>CTATTCATCGCCGTACCAGGCCTGAGCGTAGCCGCCGAAAAGACGGT<br>CTACGGCCTGAACGAATACGCGCGCATCAACGACCCGGACATCCAGC<br>TGGCCGCCAAGCTCGACACCGGCGCCAAGACCGCCTCCCTCAGCGCC<br>CGCGACATCAAGCGTTTCAAGCGCGACGGCGAAACCTGGGTGCGCTT<br>CTACCTGGCCACCGACAACGCCGACGACACCCCGATCGAGAAGCCCC<br>TGGCGCGGATCAGCAAGATCAAGCGCCGCCACGGCGACTTCAACCCC<br>GACGAGGGCAAGGCCTATACCGCCCGTCCGGTGATCGAGCTGCAGG<br>TCTGCATGGGCAAGGCGATTCGCACCATCGAAGTGAACCTGACCGAC<br>CGAAGTGCATTCCAATACCCGCTTTTGATCGGCTCGGAGGCCTTGAAG<br>AAATTCGACGCACTGGTCGATCCGAGTTTGAAATACTCGGCCGGGAAA<br>CCCGGCTGCAAACCTGATGCAAAACCTGCTGAGTCTGCGTGGTCTCAC<br>CCGCAGTTCGAAAAA**TAATCTAGAGCGGCCGCCA** | Insert for pJStrep-*rloA* |
